# The bundle sheath of rice is conditioned to play an active role in water transport as well as sulfur assimilation and jasmonic acid synthesis

**DOI:** 10.1101/2021.04.16.440137

**Authors:** Lei Hua, Sean R. Stevenson, Ivan Reyna-Llorens, Haiyan Xiong, Stanislav Kopriva, Julian M. Hibberd

## Abstract

Leaves comprise multiple cell types but our knowledge of the patterns of gene expression that underpin their functional specialization is fragmentary. Our understanding and ability to undertake rational redesign of these cells is therefore limited. We aimed to identify genes associated with the incompletely understood bundle sheath of C_3_ plants, which represents a key target associated with engineering traits such as C_4_ photosynthesis into rice. To better understand veins, bundle sheath and mesophyll cells of rice we used laser capture microdissection followed by deep sequencing. Gene expression of the mesophyll is conditioned to allow coenzyme metabolism and redox homeostasis as well as photosynthesis. In contrast, the bundle sheath is specialized in water transport, sulphur assimilation and jasmonic acid biosynthesis. Despite the small chloroplast compartment of bundle sheath cells, substantial photosynthesis gene expression was detected. These patterns of gene expression were not associated with presence/absence of particular transcription factors in each cell type, but rather gradients in expression across the leaf. Comparative analysis with C_3_ *Arabidopsis* identified a small gene-set preferentially expressed in bundle sheath cells of both species. This included genes encoding transcription factors from fourteen orthogroups, and proteins allowing water transport, sulphate assimilation and jasmonic acid synthesis. The most parsimonious explanation for our findings is that bundle sheath cells from the last common ancestor of rice and Arabidopsis was specialized in this manner, and since the species diverged these patterns of gene expression have been maintained.

**Significance statement:** The role of bundle sheath cells in C_4_ species have been studied intensively but this is not the case in leaves that use the ancestral C_3_ pathway. Here, we show that gene expression in the bundle sheath of rice is specialized to allow sulphate and nitrate reduction, water transport and jasmonate synthesis, and comparative analysis with Arabidopsis indicates ancient roles for bundle sheath cells in water transport, sulphur and jasmonate synthesis.

## Introduction

Although the major cell types of a leaf were described in the 19th century (Haberlandt, 1884) we have an incomplete understanding of the role that each plays (Aubry et al., 2014b; Mustroph et al., 2009). This lack of knowledge hinders our understanding of how basic processes are organized but is also likely to limit the rational re-design of leaves for crop improvement. One example is associated with attempts to engineer C_4_ photosynthesis into species such as C_3_ rice to increase yields (von Caemmerer et al., 2012; Hibberd et al., 2008; Langdale, 2011). As C_4_ photosynthesis typically requires metabolic compartmentation between mesophyll and bundle sheath cells, a better understanding of these tissues in rice may facilitate such a project.

The C_4_ pathway is thought to have evolved over 60 times independently in monocotyledons and dicotyledons in response to reduced water availability and CO_2_ supply (Sage, 2004). These environmental factors reduce the ratio of carboxylation to oxygenation events at the active site of RuBisCO, and so lead to higher rates of the photorespiration (Tipple and Pagani, 2007; Sage et al., 2012). Whilst in some C_4_ species, a carbon concentrating mechanism is established within large cells (Jurić et al., 2016; von Caemmerer et al., 2014; Voznesenskaya et al., 2001) in the majority of known lineages this takes place across distinct cell-types. In these two-celled C_4_ species, RuBisCO activity is replaced with phospho*enol*pyruvate carboxylase in mesophyll cells to allow bicarbonate to be fixed into oxaloacetate. High concentrations of C_4_ acids derived from oxaloacetate build up in mesophyll cells and drive diffusion to the bundle sheath where C_4_ acid decarboxylases resupply CO_2_ to RuBisCO. This reorganization of photosynthesis is thus enabled by the presence of distinct cell types such as the mesophyll and bundle sheath (Furbank, 2016). Not only do bundle sheath cells of C_4_ plants undertake a key role in photosynthesis, but they are also the primary location of starch synthesis (Lunn and Furbank, 1997) and sulphur metabolism (Gerwick et al., 1980; Passera and Ghisi, 1982; Schmutz and Brunold, 1984; Burnell, 1984; Burgener et al., 1998). Distinct classes of bundle sheath cells have been reported in maize, with abaxial bundle sheath cells of rank-2 intermediate being specialised for phloem loading (Bezrutczyk et al., 2021). The importance of bundle sheath cells in the C_4_ leaf, and the discovery that their thickened cell walls allow them to be separated from the rest of the leaf led to them being studied intensively. In summary, in the C_4_ leaf bundle sheath cells are well characterized and carry out a variety of specialized roles.

Although the vast majority of plants use the ancestral C_3_ pathway, our understanding of gene expression in bundle sheath cells specifically, and other major cell types more generally of C_3_ leaves is poor. In *Arabidopsis thaliana* (hereafter Arabidopsis) the bundle sheath represents approximately 15% of chloroplast-containing cells in the leaf (Kinsman and Pyke, 1998). Whilst the photosynthetic apparatus is functional in C_3_ bundle sheath cells (Fryer et al., 2002; Williams et al., 1989), the absolute number of chloroplasts per cell is low. Consistent with this, reducing chlorophyll accumulation or increasing the chloroplast compartment in these cells have limited impact on leaf level photosynthesis (Janacek et al., 2009; Wang et al., 2017). Instead, it appears that the bundle sheath of Arabidopsis is specialized in sulphur metabolism and glucosinolate synthesis (Aubry et al., 2014b; Koroleva et al., 2010). Stress responsive regulation of aquaporins in bundle-sheath cells are considered important for hydraulic conductance of the whole leaf (Sade et al., 2014; Shatil-Cohen et al., 2011; Attia et al., 2020) and consistent with this, bundle sheath cells have been proposed to help maintain hydraulic integrity of the xylem (Griffiths et al., 2013; Sage, 2001) as well as regulate flux of mineral and metabolites in and out of the leaf (Leegood, 2008; Wigoda et al., 2017). In summary, compared with C_4_ plants we have a relatively poor understanding of the role of bundle sheath cells in C_3_ species, and this is particularly the case in grasses such as rice.

In roots, one approach that has been used widely to define the function of discrete cell types is to generate lines in which distinct tissues are marked with a fluorescent protein, and after protoplast isolation and cell sorting, the patterns of gene expression defined (Birnbaum et al., 2003; Brady et al., 2007). In leaves, this approach has been less widely adopted, likely due to the longer incubation times typically required to generate protoplasts, and concerns about stress and de-differentiation taking place during protoplasting (Sawers et al., 2007). Recently, rapid protoplasting followed by isolation of bundle sheath cells indicated a key role in transport (Wigoda et al., 2017), single cell RNA sequencing allowed distinct patterns of gene expression to be related to discrete cell types of the Arabidopsis leaf, and indicated that bundle sheath protoplasts were most similar to xylem cells (Kim et al., 2021).

Alternate technologies that have been applied to this problem include the isolation of ribosomes from specific cell types after they were labelled with an exogenous tag (Aubry et al., 2014b; Mustroph et al., 2009), or the use of laser capture microdissection (Jiao et al., 2009). This latter approach circumvents the need to identify promoters with highly specific expression domains and the production of transgenic lines.

Here we used an optimised laser capture microdissection protocol for RNA isolation from leaves (Hua and Hibberd 2019) to study the patterns of gene expression in bundle sheath, veinal (including phloem and xylem parenchyna, xylem, as well as sieve elements and companion cells), and mesophyll cells of rice. We had three main hypotheses. First, that as the rice bundle sheath contains a small chloroplast compartment, gene expression would be poorly set up for photosynthesis. Second, as in other species, gene expression in the rice bundle sheath would favour water transport. Third, in contrast to Arabidopsis, as rice does not make glucosinolates there has been no selection pressure to restrict sulphur assimilation to the bundle sheath. Mesophyll and bundle sheath cells were distinguished by over-representation of terms including photosynthesis and co-enzyme metabolism in the former, and solute transport and protein synthesis in the latter. Transcripts encoding the majority of aquaporins were more abundant in bundle sheath cells, and this was also the case for transcripts encoding proteins associated with sulphur assimilation and nitrate reduction. Transcription factors that were preferential to each cell type were identified, but in most cases a gradient from veins to bundle sheath to mesophyll cells, or *vice versa*, was detected. Direct comparison to publicly available data from Arabidopsis identified groups of genes encoding aquaporins, proteins allowing sulphur assimilation, jasmonic acid synthesis and also a small number of transcription factor families that showed preferential expression in the bundle sheath of both species. Whilst these findings could be due to evolutionary convergence, the most parsimonious explanation is that the bundle sheath of the last common ancestor of monocotyledons and dicotyledons was specialised in water transport, sulphur assimilation as well as jasmonic acid synthesis, and that members of the Basic Leucine Zipper, C_2_H_2_-type Zinc Finger, DNA-binding with One Finger, Ethylene Responsive Factor, Hairy-Related Transcription-Factor, GRAS, MYB, Nuclear Factor-YB and Vascular Plant One-Zinc Finger Protein transcription factor families play ancient and conserved roles in this cell type.

## Results

### The rice bundle sheath is specialized for transport but also photosynthesis

To gain insight into the genetic basis for functional specialization associated with mesophyll, bundle sheath and veinal cells of rice, laser capture microdissection was used to isolate RNA from these tissues. Paradermal sections allowed the unambiguous identification of mesophyll cells (Figure 1A), bundle sheath cells containing large vacuoles and fewer chloroplasts (Figure 1C) and veins (Figure 1E). To isolate each cell-type with minimal cross contamination, mesophyll cells were first dissected and captured (Figure 1A, 1B). Sequential capture of bundle sheath cells (Figure 1C, 1D) followed by veinal cells was then possible (Figure 1E, 1F). RNA was extracted from each tissue and RNA Integrity Numbers (RIN) ranging from 6.0-7.3 indicated good quality. Strong peaks associated with ribosomal RNAs of the chloroplast were evident in mesophyll cells (Figure 1G), whereas in bundle sheath cells they were less abundant (Figure 1H) and in veins they were not discernable (Figure 1I).

**Figure 1.**
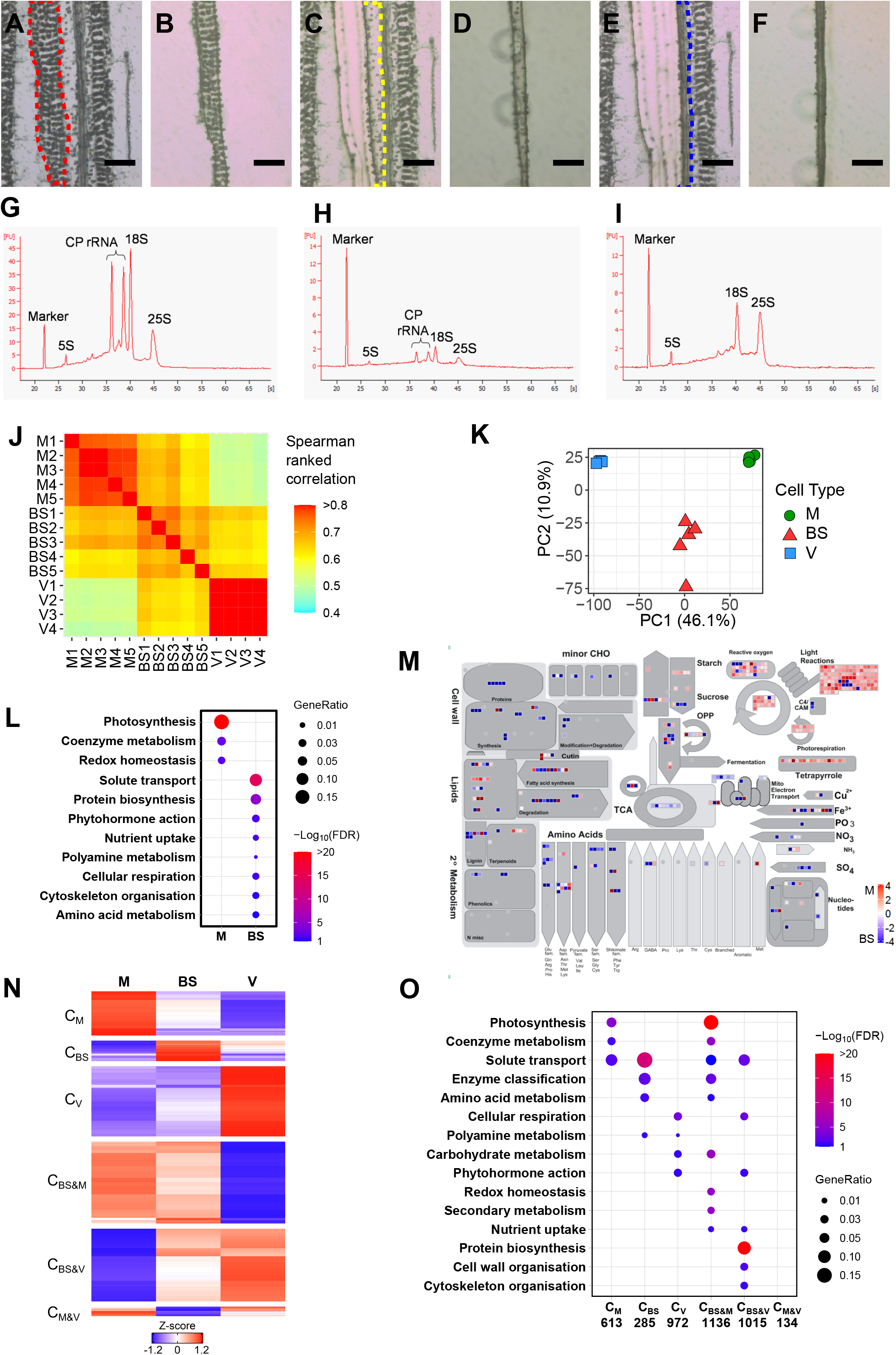
RNA was isolated from mesophyll, bundle sheath and veinal cells of rice using laser capture microdissection. Each cell-type was identified and then sequentially removed from paradermal sections prior to RNA quality being assessed. (A&B) Representative image of mesophyll cells outlined with red dash line (A) that are cut with a UV laser and then captured with an infrared laser and placed on a cap (B). (C&D) Bundle sheath cells (C) were then cut and captured (D). (E&F) Lastly, veinal cells (E) were cut and captured (F). Scale bars represents 50 microns. (G-I) Representative RNA profiles from microdissected mesophyll (G), bundle sheath (H) and veinal cells (I). Peaks from the cytosolic 25S, 18S and 5S ribosomal RNAs were detected in all cell types. Chloroplastic ribosomal RNAs (CP rRNA) were clearly detectable in mesophyll and bundle sheath cells. (J) Spearman ranked correlations of log_2_ transformed TPM (transcripts per million) indicates little variation between biological replicates from each cell type, and distinct patterns of gene expression in each cell type. (K) Principal component analysis of normalised counts after variance-stabilizing transformation showing that cell type accounted for 46.1% of the variance detected in the data. (L) Primary Mapman categories associated with differentially expressed genes in bundle sheath and mesophyll cells. Terms were defined using Fisher’s exact test (False discovery rate, FDR<0.1), colour scale represents negative log_10_ transformed FDR, gene ratio represents the ratio of matched genes in categories relative to total number of differentially expressed genes in each cell type. (M) Metabolic overviews of differentially expressed genes between bundle sheath and mesophyll. Colour scale presents the log_2_ fold change. (N) k-mean clustering of 4155 differentially expressed transcripts was performed using log_2_ transformed quantile-normalized TPM (transcripts per million) and visualized in a heatmap. Clusters were named as C_M_, C_BS_, C_V_, C_BS&M_, C_BS&V_, and C_M&V_. C_M_ contained 613 genes that were strongest in mesophyll cells, C_BS_ contained 285 genes with preferential expression in bundle sheath cells, C_V_ contained 972 genes that were strongest in veins, C_BS&M_ 1136 g.enes mostly highly expressed in mesophyll and bundle sheath cells, C_BS&V_ 1015 genes that were preferential to bundle sheath and veinal cells, and C_M&V_ 134 genes strongly expressed in both mesophyll and veinal cells. Colour scale represents Z-score. (O) Schematic illustrating the enriched primary categories derived from Mapman using Fisher’s exact test (FDR<0.1) for each of the six clusters, colour scale indicates negative log_10_ transformed FDR, size of dots (GeneRatio) represents the ratio of matched genes in each category relative to total number of genes in each cluster.

Three prime mRNA sequencing was performed and from each cell type, 24-36 million reads from four or five biological replicates obtained. After processing to remove low quality reads 13-23 million were quantified against the rice cDNA reference (MSU v7) (Supplemental Table 1). An average of 10,097, 10,083 and 13,648 transcripts were detected in each cell type (Supplemental Table 1). Spearman ranked correlation coefficients for gene expression showed little variation between replicates from each cell type, and that each cell type exhibited distinct patterns of gene expression (Figure 1J). Principle components analysis also showed close grouping of biological replicates, and that 46.1% of variance was associated with the three cell types whilst that a second component separated the bundle sheath from mesophyll and veins (Figure 1K). To assess the purity of the tissues sampled, we examined transcript abundance of genes previously reported to be associated with each cell type. Consistent with these studies, *SUCROSE PHOSPHATE SYNTHASE (SPS1*, LOC_Os01g69030) and aquaporin *PIP2;7* (LOC_Os09g36930) were preferentially expressed in mesophyll cells (Chávez-Bárcenas et al., 2000; Li et al., 2008), transcripts derived from two Tonoplast Monosaccharide Transporters *TMT1* and *TMT2* were detected in mesophyll, bundle sheath cells and vascular bundles (Cho et al., 2010), *PHOSPHOENOLPYRUVATE CARBOXYKINASE* (*PCK1*) was expressed most strongly in bundle sheath and veins (Nomura et al., 2005), and the Sucrose Transporter *SUT1* (LOC_Os03g07480) (Ibraheem et al., 2013; Scofield et al., 2007) was predominately expressed in vascular tissue (Supplemental Figure 1). The strong clustering of each cell-type combined with congruence to previous studies are consistent with the notion that these samples obtained by laser capture microdissection contained relatively little cross-contamination.

To quantify the extent to which transcript abundance differed between bundle sheath, mesophyll and veinal cells, we performed differential gene expression analysis using DESeq2 and edgeR (Love et al., 2014; Robinson et al., 2010). This identified 1,919 differentially expressed genes between bundle sheath and mesophyll cells, the majority of which (1,173) were more abundant in bundle sheath cells (FDR and adjusted *P* <5%) (Supplemental Table 2). Functional enrichment analysis identified three categories over-represented in mesophyll cells containing transcripts linked to photosynthesis, coenzyme metabolism and redox homeostasis (Figure 1L). Eight categories were associated with bundle sheath cells, including solute transport, protein biosynthesis, phytohormone action, nutrient uptake, polyamine metabolism and cellular respiration (Figure 1L). To provide an overview of metabolic specialization in mesophyll and bundle sheath cells, Mapman (Thimm et al., 2004; Schwacke et al., 2019) was used. Consistent with mesophyll cells being specialized for photosynthesis, this indicated that transcripts encoding components of the light reactions, Calvin Benson Bassham cycle and tetrapyrolle biosynthesis were upregulated in mesophyll cells, whilst those associated with many other metabolic processes including cell wall, minor carbohydrate, fatty acid, amino acid and secondary metabolism were more abundant in the bundle sheath (Figure 1M). Quantitative comparison of bundle sheath and veinal cells showed 1,258 and 660 genes were significantly up and down regulated in bundle sheath cells respectively (Supplemental Table 2) and indicated categories associated with the bundle sheath included photosynthesis, carbohydrate metabolism, redox homeostasis, secondary metabolism and solute transport (Supplemental Figure 2A). Mapman outputs confirmed transcripts encoding components of the light-dependent reactions of photosynthesis, as well as the Calvin-Benson-Bassham cycle and photorespiratory pathway were more abundant in bundle sheath than veinal cells (Supplemental Figure 2B). In contrast, transcripts preferential to veins were associated with processes that included RNA biosynthesis, protein homeostasis, lipid metabolism and solute transport (Supplemental Figure 2A). When mesophyll and veins were compared, a greater number of differentially expressed genes were identified with 1,728 and 2,038 transcripts being more abundant in mesophyll and veins respectively (Supplemental Table 2). The expected preferential expression of photosynthesis-related genes in mesophyll cells was detected, and transcripts associated with protein biosynthesis and cellular respiration were more abundant in veins (Supplemental Figure 2C&D). The greater number of differentially expressed genes between mesophyll and veins is consistent with the correlation and PCA analysis (Figure 1J&K).

To further assess patterns of transcript abundance across all three cell types we clustered genes based on expression. 4155 genes defined as being differentially expressed in the pairwise comparisons above were partitioned into six clusters associated with the cell types in which they were preferentially expressed (Figure 1N). Veins (C_V_) had the largest (972) whilst bundle sheath cells (C_BS_) had the fewest (285) number of preferentially expressed genes. Functional enrichment analysis showed that genes in the mesophyll (C_M_) were over-represented in photosynthesis, coenzyme metabolism and solute transport, whilst C_BS_ were enriched in solute transport, enzyme classification, amino acid metabolism and polyamine metabolism (Figure 1O). Genes in C_V_ were involved in cellular respiration, polyamine metabolism, carbohydrate metabolism and phytohormone action (Figure 1O). C_BS&M_ contained genes highly expressed in both mesophyll and bundle sheath, and was over-represented in processes including photosynthesis, coenzyme metabolism, carbohydrate metabolism, redox homeostasis, secondary metabolism (Figure 1O). C_BS&V_ contained genes associated with protein biosynthesis, cellular respiration, and solute transport (Figure 1O). No enriched categories were associated with both the mesophyll and vein cells (C_M&V_). Consistent with their distinct function, vein and mesophyll clusters showed low overlap, but the most abundant transcripts in bundle sheath cells were also either expressed in veins or mesophyll cells. Overall, and associated with their morphology, the data reveal a gradient in photosynthesis-related transcripts from low in veins to high in mesophyll cells.

### Patterning of gene expression in the rice bundle sheath conditions the cells for water transport

To understand the gene classes responsible for enrichment of the transport term in the bundle sheath, we examined expression of major transporter families in each of the six clusters. The family most enriched in C_BS_ was the major intrinsic protein (MIP) group, but multiple genes belonging to the cation diffusion facilitators (CDF) superfamily, major facilitator superfamily (MFS), ion transporter (IT) superfamily, multidrug/oligosaccharidyl-lipid/polysaccharide (MOP) flippase superfamily and amino acid-polyamine-organocation (APC) superfamily were also present (Supplemental Figure 3A).

The MIP group contains genes encoding water channels (aquaporins) including plasma membrane intrinsic proteins (PIPs), tonoplast intrinsic proteins (TIPs) (Sakurai et al., 2005). Transcripts encoding three and two members of the PIP1 and PIP2 families respectively accumulated preferentially in the bundle sheath compared with mesophyll and veinal cells (Figure 2A). Consistent with the deep sequencing data, *in situ* RNA localization of *PIP1*.*1* and *PIP1*.*3* generated strong signal associated with the periphery of bundle sheath cells (Figure 2B). Transcripts encoding three tonoplast intrinsic proteins (*TIP1*.*1, TIP1*.*2* and *TIP2*.*2*) also accumulated preferentially in bundle sheath cells (Figure 2A), presumably allowing water storage in the large vacuole. Lastly, in addition to these transporters, specific P-type and V-type ATPases that could regulate leaf hydraulic conductance (Grunwald et al., 2021) and establish a proton gradient across plasma and vacuole membranes to power secondary transport were strongly expressed in bundle sheath cells (Supplemental Figure 3B). Taken together, this preferential patterning of multiple aquaporins to the rice bundle sheath suggests an important role for these cells associated with water transport and storage.

**Figure 2.**
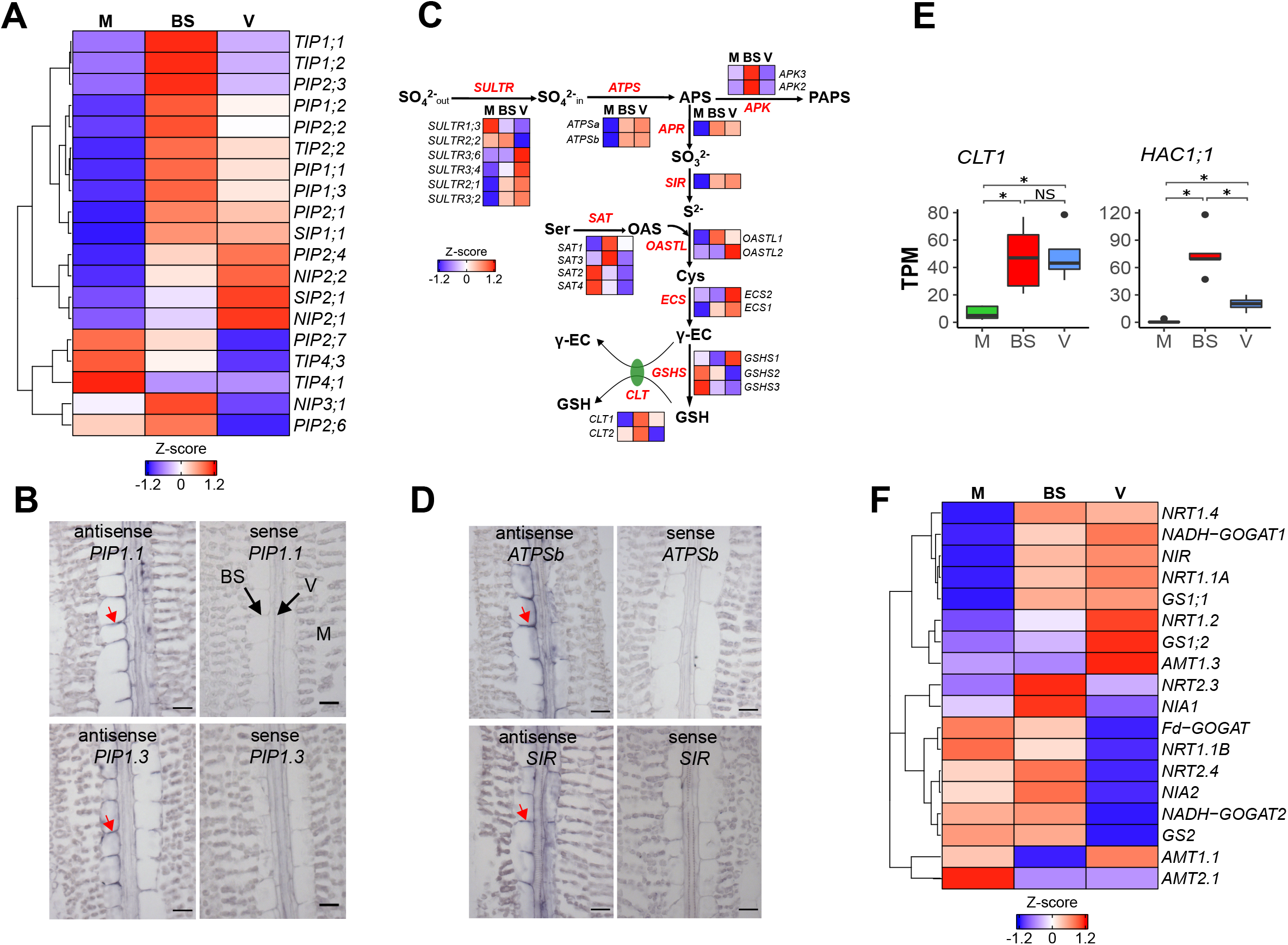
Preferential accumulation of transcripts associated with water transport, sulphur and nitrogen assimilation in the rice bundle sheath. (A) Relative transcript abundance for aquaporins in mesophyll, bundle sheath and veinal cells. Log_2_ transformed quantile normalized TPM were scaled and genes with similar expression pattern were clustered using hierarchical method. (B) Representative image after *in situ* hybridization localization for *PIP1*.*1* and *PIP1*.*3* mRNAs, scale bars represent 20 microns, red arrows indicate specific signal on bundle sheath cell periphery. (C) Schematic illustrating sulphur assimilation and relative transcript abundance in mesophyll (M), bundle sheath (BS) and veinal (V) cells depicted as Z-score from log_2_ transformed quantile normalized TPM. (D) Representative image after *in situ* hybridization for *ATPSb* and *SIR* mRNAs, scale bars represent 20 microns, red arrows indicate specific signal on bundle sheath cell periphery. (E) Transcript abundance of *CLT1* and *HAC1;1*. (F) Relative transcript abundance for genes involved in nitrogen assimilation. Abbreviations for (A): PIP, plasma membrane intrinsic proteins; TIP, tonoplast intrinsic proteins; SIP, small basic intrinsic proteins; NIP, NOD26-like intrinsic proteins. Abbreviations for (C): SULTR, sulphate transporter; ATPS, ATP sulfurylase; APS, adenosine 5’-phosphosulfate; PAPS, 3’-phosphoadenosine 5’-phosphosulfate; APR, APS reductase; SIR, sulfite reductase; APK, APS kinase; SAT, serine acetyltransferase; OAS, O-acetylserine; OASTL, O-acetylserine (thiol)lyase; γ-ECS, glutamate-cysteine ligase; γ-EC, γ-glutamylcysteine; CLT, chloroquine-resistance transporter-like transporter; GSHS, glutathione synthetase. Abbreviations for (F): NRT, nitrate transporter; AMT, ammonia transporter; NIA, nitrate reductase; NIR, nitrite reductase; GS, glutamine synthetase; NADH-GOGAT, NADH-dependent glutamate synthase; Fd-GOGAT, ferredoxin-dependent glutamate synthase.

### The rice bundle sheath preferentially accumulates transcripts associated with sulphur and nitrogen assimilation

Sulphur is an essential element required for both central metabolism and responding to biotic and abiotic stress. Although primarily taken up as sulphate by roots, reduction mainly takes place in leaves (Figure 2C). Prior to activation by ATP sulfurylase (ATPS) to adenosine 5’-phosphosulfate (APS), transport into the cell is mediated by SULTR transporters. APS is reduced into sulfite and sulfide by APS reductase (APR) and sulfite reductase (SIR) respectively, and then incorporated into O-acetylserine via O-acetylserine (thiol)lyase (OASTL) to generate the amino acid cysteine. Notably, transcripts derived from two highly expressed *SULTR* transporters (*SULTR2;1, SULTR3;2*), both *ATPS* genes, *APR*, as well as *SIR* and *OASTL1* were more abundant in bundle sheath cells compared with the mesophyll (Figure 2C). With the exception of transcripts encoding *APR* and *OASTL1* that were most abundant in bundle sheath cells, most were even more highly expressed in veins (Figure 2C). RNA *in situ* hybridization for transcripts encoding ATPS and SIR showed stronger signals in bundle sheath and veins compared with mesophyll cells (Figure 2D). These results strongly imply that gene expression of the rice bundle sheath cells is conditioned to allow sulphur assimilation.

The data also indicate that gene expression in the bundle sheath is conditioned for synthesis of glutathione, a major sulphur-containing metabolite that plays critical roles in redox homeostasis and heavy metal(loid) detoxification. Biosynthesis of glutathione is catalyzed by γ-glutamylcysteine synthetase (ECS) to generate γ-glutamylcysteine (γ-EC) from glutamate and cysteine, followed by ligation of glycine and γ-EC by glutathione synthetase (Foyer and Noctor, 2011; Hernández et al., 2015). The intermediate γ-EC has to be exported from the plastid by the CRT-like transporter (CLT) (Maughan et al., 2010; Hernández et al., 2015; Yang et al., 2016) to sustain GSH biosynthesis in the cytosol (Pasternak et al., 2007). Interestingly, we found that transcripts encoding ECS1 were preferentially expressed in bundle sheath and veinal cells (Figure 2C), and that γ-glutamylcysteine transporter *CLT1* accumulated preferentially in bundle sheath cells (Figure 2C). Rice absorbs both arsenate and arsenite by different transporters, but arsenate needs to be reduced into arsenite before it can be detoxified by phytochelatin. *CLT1* has been reported to be critical for rice tolerance to arsenic because it determines phytochelatin biosynthesis and arsenate reduction (Yang et al., 2016). Notably, the arsenate reductase *HAC1;1* also showed preferential expression in the bundle sheath, suggesting that this cell type may play an important role in arsenate reduction and detoxification (Figure 2E).

As with sulphur assimilation, transcripts encoding some of the pathway allowing nitrate reduction were more highly expressed in the bundle sheath and veinal cells compared with mesophyll cells. Interestingly, this included the nitrate transporters *NRT1*.*4, NRT1*.*1A, NRT1*.*2*, and *NRT2*.*3*, both nitrate reductases (*NIA1* and *NIA2*), nitrite reductase (*NIR*) and glutamine synthetase (*GS1*.*1*). Transcripts encoding *NRT2*.*3, NIA1* and *NIA2* were most abundant in bundle sheath cells, while the rest were also highly expressed in veins. In contrast, transcripts encoding glutamine synthetase (*GS2*) and glutamate synthase (*Fd-GOGAT*) that allow ammonia assimilation in the chloroplast were preferentially expressed in the bundle sheath and mesophyll relative to veinal cells (Figure 2F, Supplemental Figure 4). These results indicate that gene expression in the rice bundle sheath is also tuned to specialise in nitrate assimilation and amino acid biosynthesis.

### The Calvin Benson Bassham cycle, photorespiration and C_4_ cycle gene expression

Compared with the mesophyll, bundle sheath cells contain few chloroplasts, and veins only contain rudimentary plastids (Figure 3A). Consistent with this, transcripts encoding most components of photosynthetic electron transport chain were abundant in mesophyll cells, and whilst barely detectable in veins they were clearly expressed in the bundle sheath (Supplemental Figure 5A). Exceptions included one ferredoxin that showed highest transcript abundance in bundle sheath cells, three ferredoxin genes for which transcripts were more abundant in bundle sheath and vein cells compared with the mesophyll, and three homologs of photosystem I subunit PsaA, photosystem II subunit PsbK, and (Supplemental Figure 5A, 5B). In contrast to the primary ferredoxin Fd1 (LOC_Os08g01380) involved in photosynthetic electron transport (He et al., 2020) and primarily expressed in mesophyll cells, four other ferredoxins including the nitrate-inducible Fd (LOC_Os05g37140) (Doyama et al., 1998) were preferentially expressed in in bundle sheath and veins indicating that rice bundle sheath cells have optimised reducing power for nitrate reduction, as well as sulphur assimilation. A gradient from high expression in mesophyll to low in veins was observed for most enzymes of the Calvin Benson Bassham cycle. Exceptions included *RUBISCO ACTIVASE* (*RCA*), which was poorly expressed in both bundle sheath and veinal cells, and RBCS4 which had similar transcript abundance in bundle sheath and mesophyll cells (Figure 3B, Supplemental Figure 5C). The ratio of *RCA* to *RbcS* transcripts was twofold higher in mesophyll compared with bundle sheath cells. With the exception of ADP-glucose pyrophosphorylase subunits (APL1, APS2), starch synthase (SSI) and granule-bound starch synthase (GBSSII) which were more abundant in mesophyll cells, transcripts encoding enzymes of starch synthesis were similar in mesophyll and bundle sheath cells (Supplemental Figure 5D). Although transcripts associated the glucose 6-phosphate/phosphate translocator (GPT1, GPT2-1) were more abundant in bundle sheath than mesophyll cells, their abundance was low compared with the triose phosphate/phosphate translocator (TPT1) (Supplemental Figure 5D). Together, these data indicate that gene expression in the bundle sheath is set up to favour starch synthesis using products of the Calvin Benson Bassham cycle.

**Figure 3.**
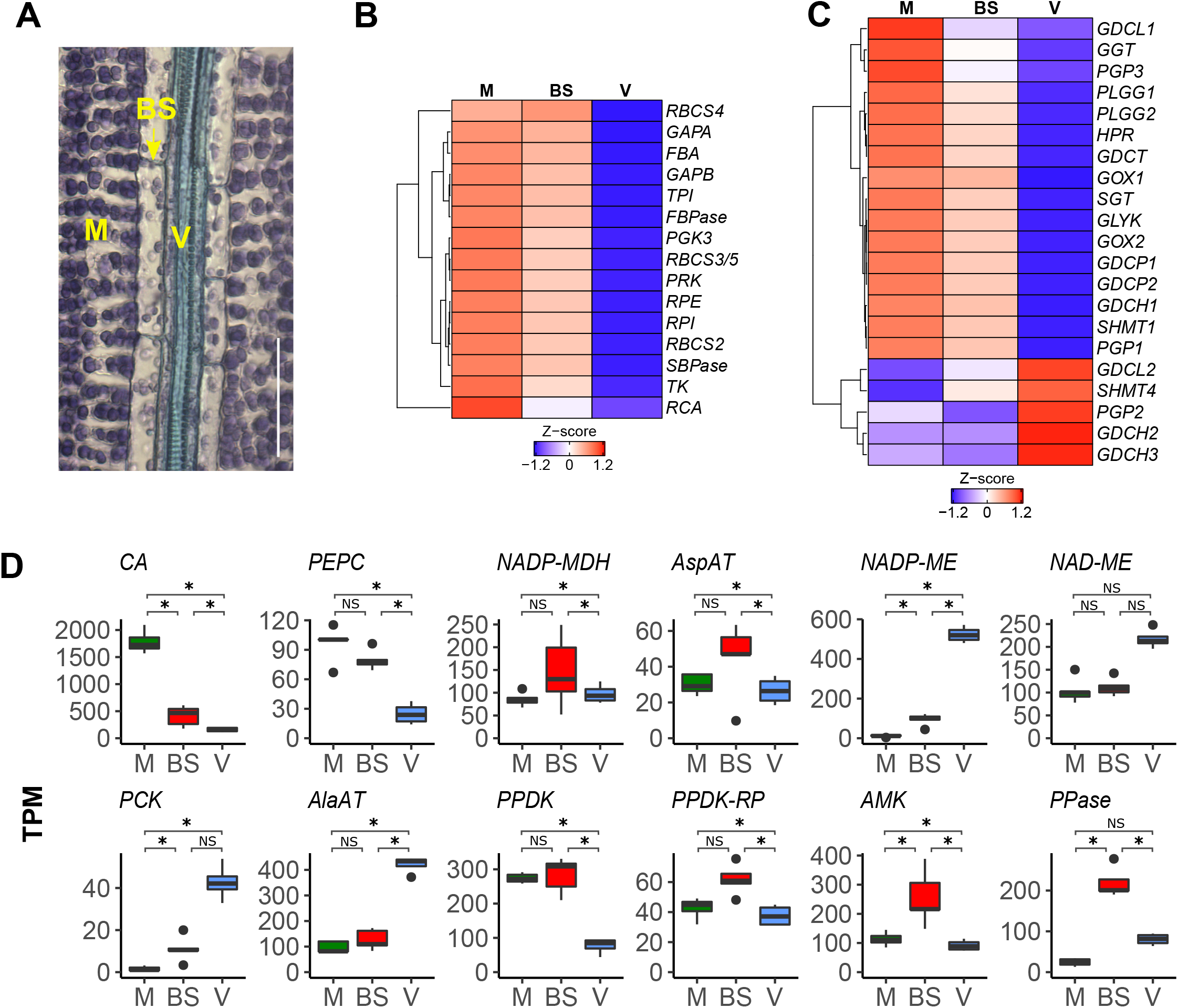
Relative transcript abundance for genes involved in Calvin Benson Bassham cycle (B) and photorespiration (C) and C_4_ pathway (D) in mesophyll (M), bundle sheath (BS) and veinal (V) cells of rice. (A) Paradermal section of rice leaf stained with toluidine blue shows bundle sheath cells are less occupied by chloroplasts compared with mesophyll cells, scale bar represents 50 microns. (B&C) Transcripts associated with Calvin Benson Bassham cycle (B) and photorespiration (C) were preferentially expressed in mesophyll cells, log_2_ transformed quantile normalized TPM were scaled and genes with similar expression pattern were clustered using hierarchical method. (D) Transcripts encoding the C_4_ acid decarboxylases NADP-DEPENDENT MALIC ENZYME (NADP-ME) and PHOSPHOENOLPYRUVATE CARBOXYKINASE (PCK) accumulated preferentially in bundle sheath and veinal cells whilst ancillary enzymes AMP KINASE (AMK) and PYROPHOSPHORYLASE (PPASE) for PYRUVATE,ORTHOPHOSPHATE DIKINASE (PPDK) accumulated preferentially in bundle sheath cells, data are presented as TPM (transcript per million), asterisks indicate statistically significant difference (FDR and adjust *P* < 0.05 using edgeR and DESeq2 analysis) Abbreviations in (B): PGK, phosphoglycerate kinase; GAPA/B, glyceraldehyde-3–phosphate dehydrogenase; TPI, triose-phosphate isomerase; FBA, fructose-1,6-bisphosphate aldolase, FBPase, fructose-1,6–bisphosphatase; TK, transketolase; SBPase, sedoheptulose 1,7–bisphosphatase; RPI, phosphopentose isomerase; RPE, phosphopentose epimerase; PRK, phosphoribulokinase; RBCS, RuBisCO small subunit; RCA, RuBisCO acvase. Abbreviations in (C): PGP, phosphoglycolate phosphatase; GOX, glycolate oxidase; GGT, glutamate:glyoxylate aminotransferase; GDC, glycine decarboxylase; SHMT, serine hydroxymethyltransferase; SGT, serine:glyoxylate aminotransferase; HPR, hydroxypyruvate reductase; GLYK, glycerate kinase. Abbreviations in (D): CA, carbonic anhydrase; PEPC, phospho*enol*pyruvate carboxylase; NADP-MDH, NADP-dependent malic dehydrogenase; AspAT, aspartate aminotransferase ; NADP-ME, NADP-dependent malic enzyme; NAD-ME, NAD-dependent malic enzyme; PCK, phospho*enol*pyruvate carboxykinase; AlaAT, alanine aminotransferase; PPDK, pyruvate,orthophosphate dikinase; PPDK-RP, regulatory protein of PPDK; AMK, AMP kinase; PPase, pyrophosphorylase.

Most transcripts encoding proteins of photorespiration showed strong expression in the mesophyll, weaker expression in the bundle sheath and very poor expression in veins (Figure 3C, Supplemental Figure 5E). The only exceptions were genes with very low absolute levels of expression that were preferential to veins, and which included one L-*GLYCINE DECARBOXYLASE* subunit (*GDCL2*), one *SERINE HYDROXYMETHYLTRANSFERASES* (*SHMT4*), one *PHOSPHOGLYCOLATE PHOSPHATASE* (*PGP2*) and two *GLYCINE DECARBOXYLASE COMPLEX H* subunits (*GDCH2*&*3*) (Figure 3C, Supplemental Figure 5E).

It has previously been reported that vascular tissue of C_3_ plants possesses high activities of some enzymes associated with C_4_ photosynthesis, including all three C_4_ acid decarboxylases and pyruvate,orthophosphate dikinase (Hibberd and Quick, 2002; Brown et al., 2010; Shen et al., 2016). However, to our knowledge it has not been possible to delimit these activities to the specific cells associated with the vascular tissue. To investigate this, we assessed abundance of transcripts encoding core enzymes of the C_4_ cycle in veins, bundle sheath and mesophyll cells of rice (Figure 3D). Transcripts of *CARBONIC ANHYDRASE* (*CA*) were more abundant in the mesophyll compared with bundle sheath and vein cells, a pattern consistent with that required for C_4_ photosynthesis. Transcripts encoding PEP carboxylase (PEPC), NADP-MALIC DEHYDROGENASE (NADP-MDH), ASPARTATE AMINOTRANSFERASE (AspAT), NAD-MALIC ENZYME (NAD-ME), ALANINE AMINOTRANSFERASE (AlaAT), PYRUVATE,ORTHOPHOSPHATE DIKINASE (PPDK) and the PPDK REGULATORY PROTEIN (PPDK-RP) showed no significant difference in abundance between mesophyll and bundle sheath cells. However, transcripts encoding AMP KINASE (AMK) and PYROPHOSPHORYLASE (PPASE), which allow the PPDK reaction and APS synthesis by ATP sulfurylase to proceed, showed higher abundance in the bundle sheath than both veins and mesophyll cells (Figure 3D), suggesting that the activity of PPDK might be higher in this cell type. Notably, two C_4_ acid decarboxylases *NADP-DEPENDENT MALIC ENZYME* (*NADP-ME*) and *PHOSPHOENOLPYRUVATE CARBOXYKINASE* (*PCK*) showed a gradient in transcript abundance from veins, to bundle sheath to mesophyll cells. These data indicate that high activity of these C_4_ decarboxylases in vascular bundles of rice (Shen et al., 2016) is likely caused by their expression in the vein rather than bundle sheath cells. Overall, C_4_ genes could be partitioned into three main groups, either showing a strong negative gradient (*CA* and *PEPC*) from mesophyll to vein cells, a strong positive gradient (*NADP-ME, NAD-ME, PCK* and *AlaAT*) or a tendency to be most strongly expressed in the bundle sheath (*NADP-MDH, AspAT, PPDK, PPDK-RP, AMP* and *PPase*) (Figure 3D). This finding is consistent with them being part of multiple gene regulatory networks in the ancestral C_3_ state that pattern their expression across these cell types.

### Transcription factors and their cognate *cis*-elements associated with the rice mesophyll, bundle sheath and vein

To gain insight into the regulatory architecture associated with the three cell types, we identified transcription factors in each of the six clusters and designated these as TF_M_, TF_BS_, TF_V_, TF_BS&M_, TF_BS&V_ and TF_M&V_. From a total of 201 differentially expressed transcription factors, over 30% of them (66) showed vein-specific expression (TF_V_, Figure 4A), including families such as bZIP, bHLH, G2-like, MYB-related and Dof (Supplemental Figure 6A). ERF, HD-ZIP and MYB-related were the most abundant transcription factor families in mesophyll-specific cluster TF_M_ (Figure 4A, Supplemental Figure 6A). Transcription factors known to regulate chloroplast biogenesis and photosynthesis were most abundant in mesophyll cells, but also detected in the bundle sheath. This included GNC (LOC_Os06g37450) in TF_M_, and GLK1 (LOC_Os06g24070), GLK2 (LOC_Os01g13740) and CGA1 (LOC_Os02g12790) in TF_BS&M_. Thus, consistent with the chloroplast complement of each cell type (Figure 3A), these transcription factors found in TF_BS&M_ were less abundant in bundle sheath compared with mesophyll cells (Supplemental Figure 6C).

**Figure 4.**
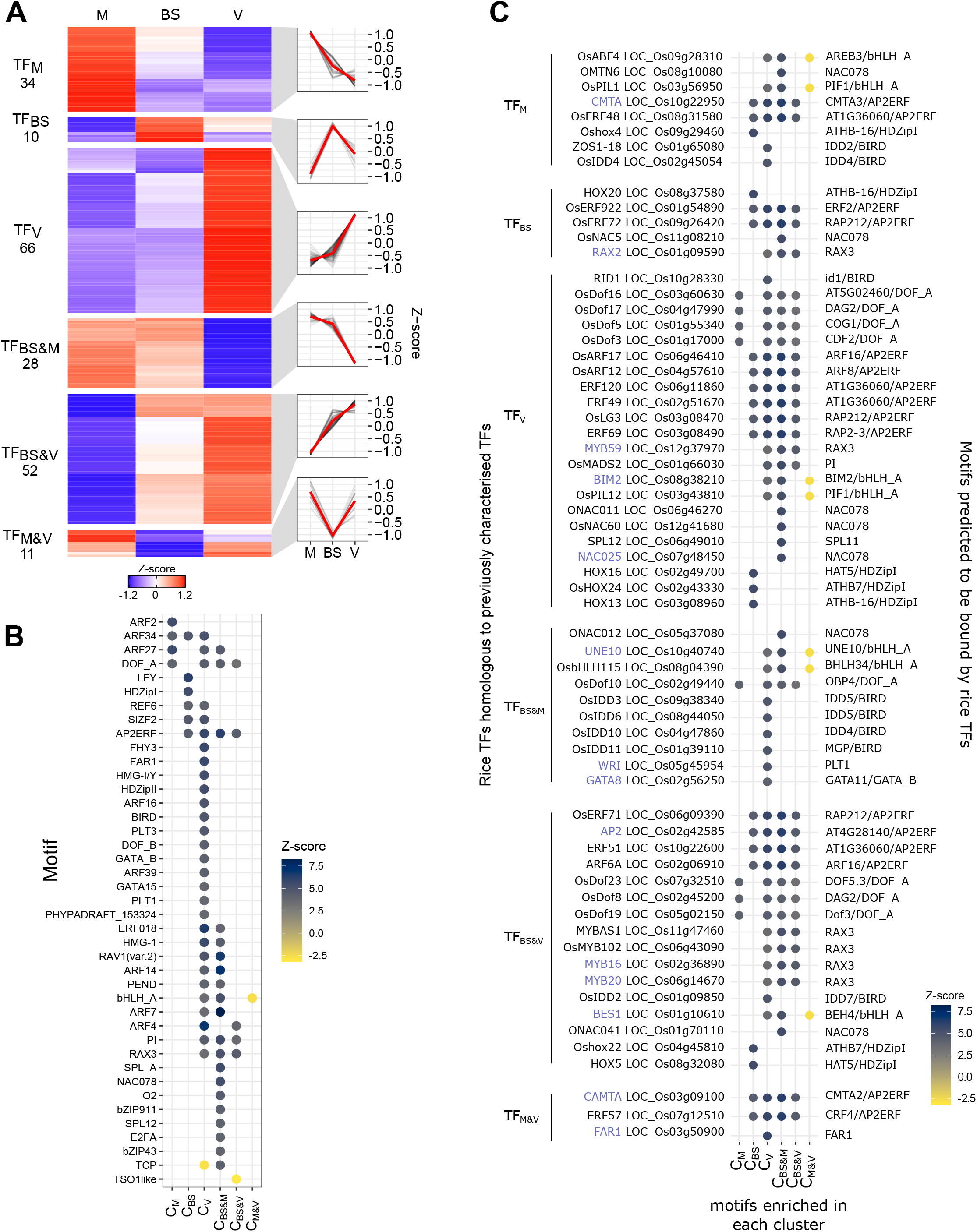
Patterning of transcription factors between mesophyll, bundle sheath and veinal cells of rice. **(A)** Transcription factors from cluster C_M_, C_BS_, C_V_, C_BS&M_, C_BS&V_, and C_M&V_ were designated as TF_M_, TF_BS_, TF_V_, TF_BS&M_, TF_BS&V_, and TF_M&V_ respectively, relative abundance of differentially expressed transcription factors were presented as heatmap and line plot of Z-score which is calculated from log_2_ transformed quantile normalized TPM, red lines in line plot represent mean of Z-score. **(B)** Significantly enriched or depleted motifs were identified in each of the cistromes from the 6 gene expression clusters, enrichment was calculated using the regioneR permutation testing package (Gel et al., 2016) following motif scanning using FIMO to identify motifs from the plant Jaspar non-redundant database (Fornes et al., 2020). The Z-scores are shown with a colour scale to show the magnitude of enrichment (dark blue) or depletion (yellow) for motifs that were significant aer multiple testing correction. Motifs derived from closely related TFs were grouped together for visualisaon based on their degree of overlap to predicted target sites (e.g. AP2ERFs). The cistrome from cluster C_V_ shows the greatest number of enriched motifs, including 13 uniquely enriched, while the C_M_ and C_BS_ cistromes have far fewer.**(C)** Cluster specific TFs (left of panel) were mapped to motifs (right of panel) they would be most likely to bind based on high protein sequence similarity with the proteins in the Jaspar plant motif database. The TFs that mapped to any enriched motifs are shown with the motif enrichment data. This allows visualisation of the intersection between TF transcript abundance with potential acvation activity. Gene symbols of rice transcripon factors were retrieved from funRiceGene database (Yao et al., 2018) but for the symbols not found in the database, symbols of best hit Arabidopsis transcription factors were used and presented in blue. The matching motifs show first the best match and then the motif group if part of a group as shown in 4B.

The bundle sheath specific cluster contained only ten genes that derived from families such as the ERFs, bZIPs and bHLHs (Figure 4A, Supplemental Figure 6A). Transcription factors abundant in both bundle sheath and vein belonged to the MYB, ERF, bZIP and WRKY families, whilst CO-like, C_2_H_2_, and G2-like families were abundant in the mesophyll and bundle sheath cluster (Figure 4A, Supplemental Figure 6A). The ZF-HD, G2-like, DBB, MIKC-MADS, and Dof families were significantly enriched in the vein-specific cluster, CO-like transcription factors were over-represented in TF_BS&M_, and Dof, MYB, HRT-like and VOZ transcription factors were over-represented in TF_BS&V_ (Supplemental Figure 6B).

To determine whether DNA motifs known to be bound by transcription factors were associated with genes preferential to each cell type, we performed motif enrichment analyses using the FIMO and AME tools (Bailey et al., 2015). Although the two approaches differ in their statistical testing they returned broadly similar estimates for the number of enriched motifs. AME found the highest number of motifs in C_BS_ and C_BS&M_ (n=44 and 91 respectively), whilst FIMO identified the most enriched motifs in C_V_ and C_BS&M_ (n=86 and 100 respectively) (Supplemental table 4 and 5). Both found either few, or no enriched motifs in the C_BS&V_ (AME=11; FIMO=5), C_M_ (AME=0; FIMO=3) or C_M&V_ (both 0) clusters (Supplemental table 4) suggesting that genes making up these three clusters are heterogeneously regulated.

As many DNA motifs share considerable sequence similarity, and closely related transcription factors can bind to similar motifs, we collapsed known DNA binding motifs from closely related transcription factors into forty-one groups (Figure 4B). Twenty-three of these groups were enriched or depleted in a specific cluster associated with transcript abundance in the three cell types. Although many were found in multiple clusters, there were no examples where a particular motif was statistically enriched or depleted in all six clusters (Figure 4B). C_M_ contained the fewest enriched motifs (n=4), only one of which (ARF2) was uniquely enriched in this cluster (Figure 4B). HD-Zip I and LHY motifs were unique to C_BS_, but all other enriched motifs in the bundle sheath were also found in C_V_ suggesting overlap in the regulation of gene expression between bundle sheath and vein cells (Figure 4B). C_V_ contained the most enriched motifs (n=29) of which almost half of these were unique to this cluster and included BIRD, Dof, GATA, HD-Zip II, PLT, FAR1 and FHY3 motifs (Figure 4B). We next investigated whether transcription factors and their cognate DNA binding sites were associated with the six clusters that had been defined by patterns of gene expression across mesophyll, bundle sheath and veinal cells. As relatively few transcription factors from rice have had their DNA binding characteristics determined, we first used protein homology (BLASTP bit-score > 100) to link them with transcription factors for which DNA binding data are available. Of the mesophyll specific transcription factors, although five are likely to bind motifs enriched in the cistrome of bundle sheath and mesophyll cells (Figure 4C), none were associated with motifs enriched only in the mesophyll cistrome. In contrast, three bundle sheath specific transcription factors coincided with motifs enriched in the BS-specific cistrome. Vein-specific transcription factors showed the greatest convergence with enrichment in their cognate DNA binding sites with fifteen mapping to the vein specific cistrome. This included transcription factors predicted to bind BIRD, Dof, AP2ERF, MYB, MADS and bHLH motifs (Figure 4C). Overall, this approach identifies families of transcription factors preferentially expressed in cell types in which their cognate motifs were over-represented in the cistrome of that cell type. We regard these transcription factors as strong candidates for patterning gene expression across these cell-types.

### Conserved patterning of gene expression in bundle sheath cells from rice and Arabidopsis

Bundle sheath cells are present in both monocotyledons and dicotyledons. Previous analysis has compared transcripts loaded onto ribosomes in the bundle sheath of Arabidopsis with those from whole leaves. In so doing, it was concluded that the bundle sheath in Arabidopsis is important for sulphur and glucosinolate metabolism as well as trehalose synthesis (Aubry et al 2014b). Functional enrichment of bundle sheath and mesophyll preferential genes from rice and Arabidopsis were compared. Consistent with the mesophyll representing the major cell type conducting photosynthesis, gene sets common to the mesophyll from the two species included transcripts important for the photosynthetic electron transport chain and photophosphorylation, the Calvin Benson Bassham cycle, photorespiration, tetrapyrrolle biosynthesis, chloroplast redox homeostasis, organelle protein biosynthesis and terpenoid biosynthesis (Figure 5A). In contrast, only four categories of genes were enriched in the bundle sheath from both species, namely carrier-mediated transport, sulphur assimilation, amino acid biosynthesis and jasmonic acid action (Figure 5A).

**Figure 5.**
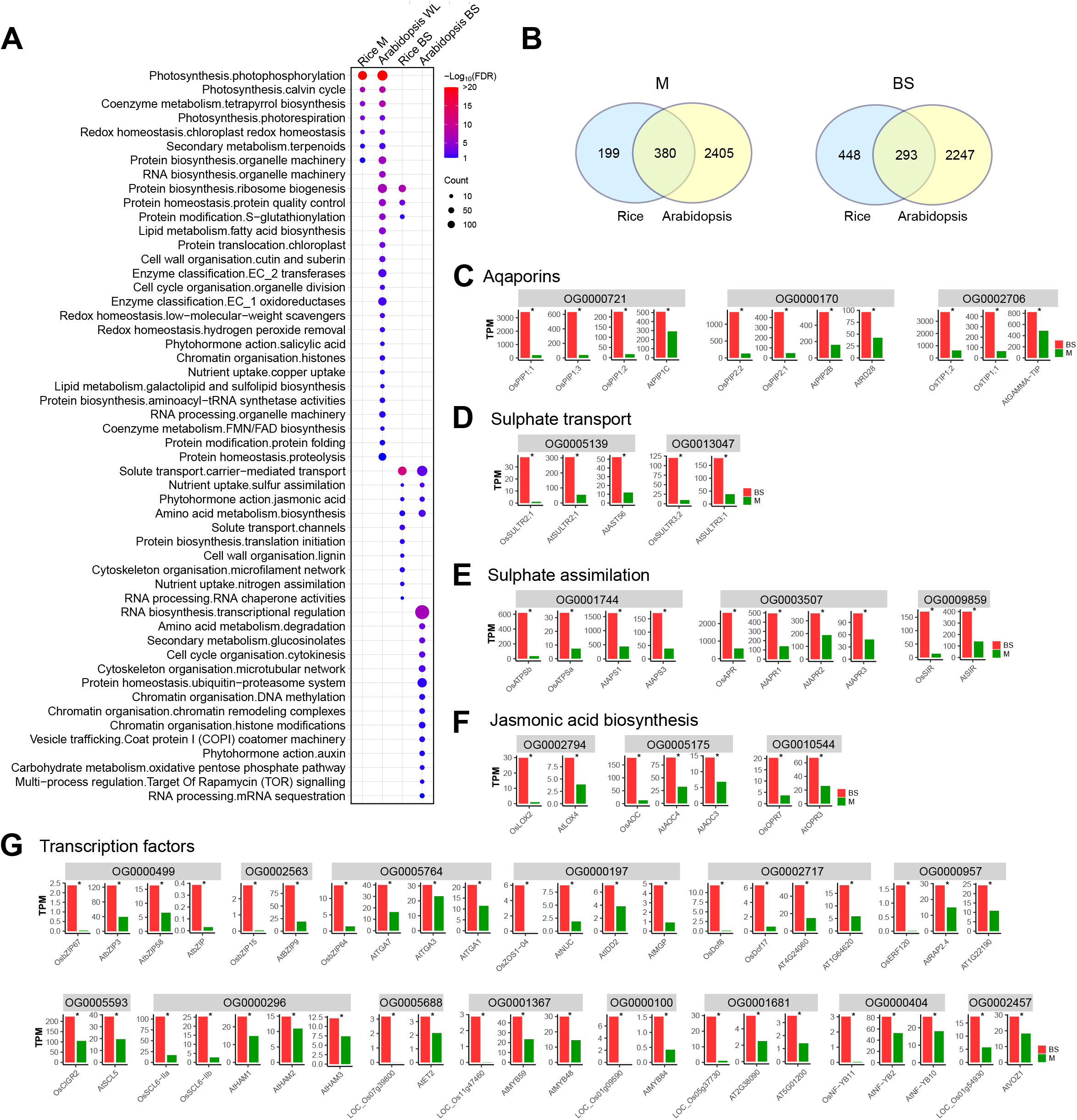
Conserved patterning of gene expression in the Arabidopsis and rice bundle sheath. Orthologs from Arabidopsis and rice associated with aquaporins, sulphate transport and assimilation as well as jasmonic acid biosynthesis are strongly expressed in bundle sheath cells. (A) The enriched Mapman categories (secondary level) of BS and M preferential genes in rice and Arabidopsis were defined using Fisher’s exact test (FDR<0.1). (B) Venn diagram illustrating the extent to which genes in the same orthogroup are preferentially expressed in mesophyll or bundle sheath cells of both rice and Arabidopsis. (C-G) Transcript abundance of Arabidopsis and rice genes belonging to the same orthogroups of aqaporins (C), sulphate transport (D), sulphate assimilation (E), jasmonic acid biosynthesis (F), and transcription factors (G). Data are presented as TPM and statistically significant differences annotated with an asterisk (FDR and adjust *P* < 0.05 using edgeR and DESeq2 analysis in this study, PPDE>0.95 in Aubry et al., 2014b), red and green bars represent bundle sheath and mesophyll respectively.

We next identified genes from both species likely to be descended from a common ancestor and placed these into orthogroups (Emms and Kelly, 2019). Genes were assigned to 10,665 orthogroups and the extent to which each was preferentially expressed in mesophyll or bundle sheath cells determined. Genes from 380 orthogroups were preferentially expressed in mesophyll cells of rice and whole leaves of Arabidopsis, whilst genes from 293 orthogroups were shared by bundle sheath of both species (Figure 5B), in both cases a greater overlap than would be expected by chance (Fisher’s exact test, *p*< 2.2e^-16^ for both bundle sheath and mesophyll). However, the odds ratio that is also generated from the Fisher’s exact test indicated that there was a greater degree of overlap between the mesophyll of rice and whole leaves of Arabidopsis than between their bundle sheath cells (odds ratios were 5.32 and 2.06 for mesophyll and bundle sheath respectively). We thus conclude that the expression of orthologs has diverged more in the bundle sheath than the mesophyll of these species. Specific orthogroups associated with the mesophyll in both species were mainly associated with photosynthesis but also included eleven orthogroups of transcription factors, one of which included the transcription factor *GLK1* (OG0002393) (Supplemental table 5) that is known to activate photosynthesis related gene expression (Waters et al., 2008; Waters et al., 2009). Although fewer orthogroups were common to the bundle sheath in rice and Arabidopsis, overlap included three groups encoding aquaporins (PIP1 - OG0000721, PIP2 -OG0000170 and TIP1 - OG0002706), as well as orthologs involved in sulphate transport (OG0005139 and OG0013047), sulphate assimilation (OG0001744, OG0003507 and OG0009859) and jasmonic acid biosynthesis (OG0002794, OG0005175, OG0010544) (Figure 5C-F, Supplemental Figure 7, Supplemental Table 5). We compared bundle sheath preferentially expressed genes of rice and Arabidopsis with the bundle sheath marker genes identified through single cell sequencing of Arabidopsis (Kim et al., 2021). As sequencing depth from single celled sequencing is not as great, fewer transcripts were detected (Supplemental Figure 8A). However, 41 orthogroups were identified containing at least one gene preferential to the bundle sheath in all three studies, including two sulphate transporters (OG0005139 and OG0013047), ATP sulfurylase (OG0001744), and allene oxide cyclase (OG0005175). To investigate whether these genes are associated with bundle sheath cells in a broader range of species, we assessed transcript abundance of consensus orthogroups associated with water transport, sulphur assimilation, jasmonic acid biosynthesis as well as nitrate reduction in publicly available data from *Panicum virgatum* (Rao et al., 2016), *Sorghum bicolor* (Emms et al., 2016), *Setaria italica* (John et al., 2014) and *Zea mays* (Chang et al., 2012) as well as the C_4_ dicotyledon *Gynandropsis gynandra* (Aubry et al., 2014a). Except for nitrate reduction genes that are more highly expressed in mesophyll cells, this indicated that in the majority of these species, genes in each of orthogroup are preferentially expressed in bundle sheath cells (Supplemental Figure 9). Taken together, the data strongly suggest that sulphur assimilation, jasmonic acid biosynthesis, and water transport represent ancestral functions associated with the bundle sheath derived from the last common ancestor of monocotyledons and dicotyledons.

We next wished to identify if any transcription factors were preferentially expressed in bundle sheath cells of both species. The rice ortholog of the key regulator of the sulphur starvation response *SULFUR LIMITATION1* (Maruyama-Nakashita et al., 2006), *OsEIL3* (LOC_Os09g31400) showed low expression, but consistent with Arabidopsis, was more strongly expressed in bundle sheath compared with mesophyll cells (Supplemental Figure 7). Fourteen transcription factor orthogroups were identified containing at least one ortholog in both species that was preferential to the bundle sheath (Figure 5G). These included three Basic Leucine Zipper (OG0000499, OG0002563, OG0005764), C_2_H_2_-type Zinc Finger (OG0000197), DNA-binding with One Finger (OG0002717), Ethylene Responsive Factor (OG0000957), two GRAS (OG0000296, OG0005593), Hairy-Related Transcription-Factor (OG0005688), three MYB (OG0000100, OG0001367 and OG0001681), Nuclear Factor-YB (OG0000404), and Vascular Plant One-Zinc Finger Protein (OG0002457) transcription factor families (Figure 5G, Supplemental Figure 7, Supplemental table 6). These data imply purifying selection has acted to maintain expression of these transcription factors in the bundle sheath since these species diverged from their last common ancestor prior to the divergence of the dicotyledons and monocotyledons.

## Discussion

### Patterning of photosynthesis gene expression between cell-types in the rice leaf

Separation of protoplasts followed by cell sorting has provided significant insight into gene expression in specific cells of roots (Birnbaum et al., 2003; Brady et al., 2007). In leaves, the ability to separate mesophyll and bundle sheath cells from C_4_ plants (Kanai and Edwards, 1973; Edwards and Black, 1971; Moore et al., 1984) has allowed similar levels of insight into these cells, but leaf cells from C_3_ species have been more challenging to isolate. As a consequence, although major cell-types of leaves such as the mesophyll, phloem, xylem and guard cells have well defined roles, others such as the bundle sheath are relatively poorly understood (Leegood, 2008). In Arabidopsis the bundle sheath is thought to play important roles in regulating hydraulic conductance, substrate transport and storage (Shatil-Cohen et al., 2011; Sade et al., 2014; Griffiths et al., 2013). Analysis of mRNAs resident on ribosomes showed that patterns of gene expression in the Arabidopsis bundle sheath are conditioned to facilitate sulfur metabolism and glucosinolate biosynthesis (Aubry et al., 2014b). Although, suppression of chlorophyll synthase in veinal tissue including bundle sheath cells of Arabidopsis reduced photosynthesis, growth and fitness (Janacek et al., 2009), the importance of the bundle sheath itself was not defined. Moreover, in C_3_ monocotyledons, the group containing many of our most important crops, little is known about the function of bundle sheath cells. To address this, we defined gene expression in the rice bundle sheath as well as mesophyll and vein cells using laser capture microdissection coupled with mRNA sequencing. Contrary to our expectations, this indicated that although bundle sheath cells of rice contain few chloroplasts (Wang et al., 2017; Sage and Sage, 2009), transcripts encoding components of the photosynthetic electron transport chain and the Calvin Benson Bassham cycle were clearly expressed in rice bundle sheath cells. The consequence of this finding is that expression of photosynthesis genes represents a continuum from low to medium to high in veins, bundle sheath and mesophyll cells respectively. Whether this is also the case in Arabidopsis remains to be determined. There is significant interest in activating photosynthesis the bundle sheath of rice such that mesophyll and bundle sheath cells could be engineered to carry out C_4_ photosynthesis (Wang et al., 2017; Sage, 2004). The finding that photosynthesis gene expression in the bundle sheath resembles the mesophyll more than veins indicates that it needs to be re-tuned rather than completely re-programmed to achieve this demanding aim.

### Expression of C_4_ genes in the C_3_ rice leaf

Vascular bundles of C_3_ plants are known to carry out C_4_-like metabolism via high activities of C_4_ acid decarboxylases and PPDK to make use of C_4_ acids present in the transpiration stream (Hibberd and Quick, 2002; Brown et al., 2010; Shen et al., 2016). To date, it has not been possible to define whether high activity of these C_4_ enzymes in C_3_ plants is associated with their expression in vein or bundle sheath cells. The analysis of rice we present here shows that transcripts encoding two C_4_ acid decarboxylases, NADP-ME and PCK, were more abundant in bundle sheath compared with mesophyll cells, but in both cases, transcripts were even more strongly expressed in veins. These findings suggest that the high activity of NADP-ME and PCK in vascular tissue is primarily caused by expression in veins rather than the bundle sheath. Thus, the gradient in expression of genes encoding these C_4_ acid decarboxylases from high in veins to low in mesophyll is the opposite of photosynthesis genes in these cell types. Whether this is also the case in C_3_ dicotyledons such as Arabidopsis remains to be determined.

Although there was no significant difference in *PPDK* transcript abundance between bundle sheath and mesophyll cells, we detected greater abundance of transcripts encoding AMK and PPase which carry out ancillary reactions allowing the PPDK reaction to proceed. As activity of PPDK is higher in vascular strands of rice than in mesophyll cells (Shen et al., 2016), it is therefore possible that this is caused by greater activity of AMP and PPase. PPDK is important for nitrogen recycling in Arabidopsis and tobacco (Taylor et al., 2010) and so these data also suggest that this is likely the case in rice. Although *CA* and *PEPC* transcripts in rice were most abundant in mesophyll cells, the particular isoforms these transcripts encode are predicted to generate proteins localized to the chloroplast (Masumoto et al., 2010; Chen et al., 2017). In C_4_ leaves, both of these proteins are cytosolic (Hatch, 1987; Hatch and Burnell, 1990). Overall, these results suggest that at least three modifications to expression of these genes would be required to build a C_4_ cycle into rice – amplifying the existing expression of C_4_ acid decarboxylases in bundle sheath cells, repositioning CA and PEPC proteins such that they reside in the cytosol, and expressing AMK and PPase in the mesophyll for effective PPDK activity.

### The rice bundle sheath is conditioned to allow water transport, sulphate and nitrate metabolism

Our analysis of the rice bundle sheath indicates that gene expression is poised to allow photosynthesis in this cell type. This finding is consistent with the fact that although during early leaf development plastids of the rice bundle sheath contain significant amount of starch, as the leaf matures these plastids develop into chloroplasts (Miyake, 2016). Transcripts patterns in the bundle sheath were consistent with starch being synthesized from carbon skeletons derived from the Calvin Benson Bassham cycle with strong expression of *PGI, PGM, UGP1, SSIIB* and *SSIII-1*, and low expression of hexose phosphate transporters (*GPT1, GPT2-1*) compared with the *TPT1*. Our analysis also indicates that the rice bundle sheath is patterned to facilitate water transport and storage, sulphate and nitrate assimilation. It was notable that most of highly expressed aquaporins were preferentially expressed in bundle sheath cells, and included members of the PIP1, PIP2, TIP1 and TIP2 subfamilies. It has been reported that water transport activity and plasma membrane localization of PIP1 requires interaction with PIP2 (Fetter et al., 2004; Zelazny et al., 2007) and so their co-expression in the bundle sheath is compatible with efficient water transport and storage in this cell type. Notably, analysis of publicly available data indicates that strong expression of *PIP1* and *PIP2* in the bundle sheath is also found in the C_4_ grasses *Panicum virgatum, Sorghum bicolor, Setaria italica* and *Zea mays* as well as the C_4_ dicotyledon *Gynandropsis gynandra* (Rao et al., 2016; Emms et al., 2016; John et al., 2014, Chang et al., 2012; Aubry et al., 2014a), implying that bundle sheath plays important role in water transport in the common ancestor of monocotyledons and dicotyledons (Supplemental Figure 9).

Enzymes of sulphur assimilation preferentially accumulate in the bundle sheath of C_4_ grasses and C_3_ dicotyledon Arabidopsis (Gerwick et al., 1980; Passera and Ghisi, 1982; Burnell, 1984; Schmutz and Brunold, 1984; Burgener et al., 1998; Aubry et al., 2014b; Chang et al., 2012; John et al., 2014; Emms et al. 2016; Rao et al., 2016), but this is not the case in wheat and the C_4_ dicotyledons *Flaveria* and *Gynandropsis gynandra* where ATPS and APR showed similar activity and/or transcript abundance in bundle sheath and mesophyll cells (Schmutz and Brunold, 1984; Kopriva et al., 2001; Aubry et al., 2014a). We found that transcripts associated with sulphur assimilation were restricted or preferentially localized to bundle sheath cells of rice, which is consistent with expression patterns in Arabidopsis (Aubry et al., 2014b). This was also true for genes indirectly associated with sulphur assimilation such as the PPase that increases the rate of the ATP sulphurylase reaction. Several evolutionary drivers for localization of sulphur assimilation in the bundle sheath of C_4_ grasses have been discussed including co-localisation with photorespiration as a source of serine for cysteine synthesis and protection of the reaction intermediates from oxidation by Photosystem II derived oxygen (Kopriva and Koprivova, 2005).

In Arabidopsis glucosinolate synthesis is controlled by a MYC-MYB transcription factor module that has recently been shown to pattern gene expression to the bundle sheath (Dickinson et al 2020). It seemed likely that localisation of glucosinolates to the bundle sheath of Arabidopsis led to upregulation of ATPS and APR in these cells. However, the preferential expression of genes associated with sulphur assimilation in the rice bundle sheath that does not synthesise glucosinolates indicates a much more ancient role for the bundle sheath in sulphur assimilation. It also removes the proposed connection between C_4_ photosynthesis and localisation of sulphur assimilation in the bundle sheath. More broadly, it appears that the bundle sheath is conditioned for sulphur assimilation in all species analysed to date. In plants such as wheat, *Flaveria* and *Gynandropsis*, mesophyll cells are also used (Schmutz and Brunold, 1984; Koprivova et al., 2001; Aubry et al., 2014a), but in others the pathway appears to be restricted to the bundle sheath (Supplemental Figure 9). It is not clear whether the ancestral state was for gene expression allowing sulphur assimilation to be associated with bundle sheath and mesophyll cells or whether in some species expression in mesophyll cells has been gained. Moreover, a full complement of transporters and enzymes for nitrate reduction were found to preferentially be expressed in rice bundle sheath cells, other enzymes that serve to provide carbon skeleton for amino acid metabolism such as PEPC (Masumoto et al., 2010) and PPDK (Taylor et al., 2010) also showed high expression in these cells. The compartmentation of nitrate and ammonia assimilation gene expression between bundle sheath and mesophyll cells of rice contrasts with their spatial separation of C_4_ grasses and *Gynandropsis gynandra*, where nitrate reduction predominately takes place in the mesophyll, and ammonia assimilation occurs in the bundle sheath (Supplemental Figure 9). The strong expression of genes associated with nitrate reduction to our knowledge has not previously been reported in any system. Furthermore, it has implications for our understanding of transitions associated with the evolution of the C_2_ and C_4_ pathways, since modelling revealed that early changes to C_2_ metabolism should induce an imbalance in nitrogen metabolism between bundle sheath and mesophyll cells (Mallmann et al., 2014). It has been proposed that this imbalance could be counteracted by upregulating genes associated with C_4_ photosynthesis (Mallmann et al., 2014). Indeed, transcripts for key genes of nitrate assimilation are consistently more strongly expressed in mesophyll cells of C_4_ plants (Supplementary Figure 9, Rao et al., 2016; Emms et al., 2016; John et al., 2014, Chang et al., 2012; Aubry et al., 2014a). The consequences and the drivers of gene expression associated with nitrate reduction being focused on the bundle sheath in rice are unknown. However, this co-localisation of sulphur and nitrate assimilation in the rice bundle sheath was associated with preferential expression of specific ferredoxins that have previously only been implicated in differences in electron transfer reactions allowing CO_2_ or mineral nutrient reduction in shoots and roots (Yonekura-Sakakibara et al., 2000).

The ancestors of rice and Arabidopsis diverged ∼140 million years ago (Chaw et al., 2004). Despite this timescale significant overlap in cell-specific gene expression in bundle sheath and mesophyll cells from these two C_3_ species was detected. We conclude that the two cell types have retained specific roles over this extended time. Less overlap was found in bundle sheath cells suggesting that the role of the bundle sheath has evolved faster than the mesophyll. However, despite these apparent changes to gene expression in the bundle sheath, some genes associated with water transport and jasmonic acid biosynthesis were preferential to bundle sheath cells in majority of species. In contrast to sulphur assimilation, we therefore propose that these processes represent ancestral functions derived from the last common ancestor of monocotyledons and dicotyledons. Moreover, a small number of transcription factors were strongly bundle sheath preferential in both species, and so we propose that these regulators underpin ancestral and conserved functions of bundle sheath cells in dicotyledons and monocotyledons.

## Materials and methods

### Plant growth condition and sample preparation

The temperate rice (*Oryza sativa* ssp. *japonica*) Kitaake was germinated and grown in a mixture of 1:1 topsoil and sand for 2 weeks in a controlled environment growth room. Temperature was set to 28°C day, 25°C night, and photoperiod at 12 hr light and 12 hr dark. Relative humidity was 60% and photon flux density 300 μmol m^−2^ s^−1^. 1cm sections from the middle of the fourth fully expanded leaves were sampled 4 hours after dawn. Leaf tissue was fixed and embedded into Steedman’s wax as described previously (Hua and Hibberd, 2019) with minor modifications. For example, rice leaves were fixed in 100% (v/v) acetone on ice for 4 hours and before embedding tissue infiltrated with 100% Steedman’s wax at 37 °C overnight. For laser capture microdissection (LCM) and RNA extraction, paradermal sections of 7 microns were prepared with a microtome and mounted on PEN membrane slides. Prior to LCM Steedman’s wax was removed by incubating slides in 100% (v/v) acetone for 1 min. LCM was performed on an Arcturus Laser Capture Microdissection platform, with isolated cells being collected on CapSure Macro Caps and RNA extracted using the PicoPure RNA Isolation Kit with on-column DNaseI treatment. RNA quality and concentration were analyzed using an Agilent Bioanalyser RNA 6000 Pico assay.

For RNA *in situ* hybridization, middle sections from the fourth fully expanded leaf were sampled and fixed overnight using FAA fixative (50% (v/v) ethanol, 5% (v/v) acetic acid and 3.7% (v/v) formaldehyde. They were then dehydrated through an ethanol series of 50%, 70%, 85%, 95% and 100% (v/v) and embedded into Steedman’s wax as described previously (Hua and Hibberd, 2019). 8 micron thick sections were obtained on a microtome and mounted onto Superfrost plus slides. Tissue pretreatment, hybridization and colour development were performed as previously described (Jackson, 1992), except that Steedman’s wax was removed by incubating slides in 100% ethanol for 5 mins twice.

### Library preparation, RNA sequencing and data processing

20 ng of bundle sheath, mesophyll and veinal RNA was used as input for the Quantseq 3’ mRNA-seq library preparation kit (Lexogen, Moll et al., 2014) according to the manufacturer’s instructions. Libraries were sequenced using Nextseq 500 sequencer to produce single-ended 150-bp reads for each sample. The leading and tailing 10-bp of Quantseq reads were trimmed and reads with a quality score less than 20 and shorter than 50-bp were removed using BBDuk (Bushnell, 2015). Transcript abundance was determined after the remaining reads were quantified using Salmon version 0.8.2 (Patro et al., 2017) against the rice cDNA reference (MSU version7), ‘--noLengthCorrection’ flag was used to disable length correction (Corley et al., 2019). Gene level abundance (TPM, transcript per million) and counts were summarized using tximport version 1.10.1 (Soneson et al., 2016) and gene level counts were used for downstream differential gene expression analysis using DESeq2 version 1.22.2 (Love et al., 2014) and edgeR version 3.24.3 (Robinson et al., 2010). Poorly expressed genes with row sum of TPM < 1 in three samples were excluded, resulting in 15168 genes; and a Benjamini-Hochberg corrected *P*-value (DESeq2) and False discovery rate (edgeR) of <0.05 were used to define differentially expressed genes.

To quantify the abundance of transcript associated with ribosomes of Arabidopsis BS and total leaf samples, reads were obtained at Sequence Read Archive (SRA, https://www.ncbi.nlm.nih.gov/sra/) Bioproject accession PRJEB5030 and quantified using Salmon version 0.8.2 against the Arabidopsis TAIR10 reference (Berardini et al., 2015) using default parameters, gene level abundance (TPM, transcript per million) was summarized using tximport version 1.10.1 (Soneson et al., 2016). Differentially expressed genes were defined according to Aubry et al., 2014b.

### Gene clustering, over-representation analysis and gene expression visualisation

Consensus differentially expressed genes identified using DESeq2 and edgeR in each pairwise comparison were used for gene expression clustering and functional enrichment analysis. Gene expression clustering were performed with K-Means method (Hartigan and Wong, 1979) using log_2_ transformed quantile normalized TPM (transcript per million). For functional enrichment, rice proteome (MSU version7) were firstly annotated by Mapman cateories using Mercator 4 (Schwacke et al., 2019), over-representation analysis of Mapman categories were performed using Fisher’s exact test and using the expressed 15168 genes as background with false discovery rate (FDR) < 0.1. Rice transcription factor families were annotated according to PlantTFDB v5.0 (Jin et al., 2017), using all expressed transcription factor as background, over-representation test were performed using Fisher’s exact test with a cutoff FDR < 0.1. Significant Mapman categories and transcription factor families were plotted using ggplot2 (Wickham, 2016). Heatmap of gene expression clusters and pathways were plotted using ComplexHeatmap package (Gu et al., 2016). Transcript abundance were presented in boxplot of TPM (transcript per million) using default settings of hinge and whisker in ggplot2.

### Motif enrichment analysis

Leaf DNase I hypersensitive sites (DHS) (Zhang et al., 2012) within 2000-bp of the genes were extracted and used as input to the AME tool (http://meme-suite.org/tools/ame, Bailey et al., 2015; McLeay and Bailey, 2010) using default settings and a custom background of all DHS regions within 2000bp of gene loci as well as for FIMO command line tool (Grant et al., 2011) using default settings. The meme format Jaspar Plant Non-Redundant Motif Database (Fornes et al., 2020, http://jaspar2018.genereg.net/downloads/) was used for both methods. FIMO scanning identified significant matches to known motifs and the frequencies of these sites were statistically tested for enrichment or depletion using permutation testing with the regioneR R package (Gel et al., 2016). For permutation testing, all DHS regions within 2000bp of genes were used as a background for random subsampling and observed frequencies were statistically tested against the random subsampling distributions. As the Jaspar motif database contains many motifs with high similarity predicted to be bound by closely related transcription factors, we grouped similar motifs together that showed a higher than 70% overlap in predicted target sites within the DHS background sequences representing motifs that were deemed highly redundant. The most strongly enriched individual motif from each such group was used as a representative value for the group and plotted using ggplot2 (Wickham and Sievert, 2016). In order to predict which *O. sativa* transcription factors might bind predicted motifs, we mapped rice transcription factors to their best match motif through protein homology with the Jaspar Non-Redundant Motif database proteins (Fornes et al., 2020). The best scoring BLASTP match was used as long as bit-score was greater than 100 to avoid spurious assignments to motifs from other TF families. For all transcription factors found in the 6 clusters, those that were mapped to an enriched motif from any of the 6 clusters was plotted using the single motif enrichment scores, rice transcription factors were annotated according to funRiceGenes database (Yao et al., 2018).

### Defining orthogroups in rice and Arabidopsis

Protein sequences of primary transcripts of *Oryza sativa* (v7.0), *Arabidopsis thaliana* (TAIR10), *Gynandropsis gynandra* (Aubry et al., 2014a), *Brachypodium distachyon* (v3.2), *Panicum virgatum* (v1.1), *Sorghum bicolor* (v1.4), *Setaria italica* (v2.1), *Zea mays* (Ensembl-18) and *Marchantia polymorpha* (v3.1) obtained from Phytozome v12.1 (Goodstein et al., 2012) were clustered into orthogroups using Orthofinder version 2.4.1 (Emms and Kelly, 2019) with the default parameters, orthogroups that contain both rice and Arabidopsis genes were used for comparative analysis in the two species. Resolved gene trees of orthogroups were visualized using R package ggtree version 2.41 (Yu, 2020).

## Code and data availability

Code associated with this manuscript and the underlying data required to generate plots are available in the Github repository: https://github.com/hibberd-lab/Hua_et_al_Kitaake_LCM. All other data are available on request.

## Accession numbers

Raw sequencing data are deposited in the National Center for Biotechnology Information under BioProject ID PRJNA702624, BioSample ID SAMN17976370 - SAMN17976383.

## Acknowledgements

The work was funded by C_4_ Rice project grant from The Bill and Melinda Gates Foundation to the University of Oxford (2015–2019), European Research Council Grant 694733 Revolution, BBSRC Grants BBP0031171 and BBL014130 to JMH. Research in SK’s lab is funded by the Deutsche Forschungsgemeinschaft (DFG) under Germany’s Excellence Strategy – EXC 2048/1 – project 390686111.

## Supplemental Data

**Supplemental Figure S1**. Transcript abundance of genes previously reported to be associated with mesophyll, bundle sheath or veins.

**Supplemental Figure S2**. Mapman categories and metabolic overview of pairwise comparisons between bundle sheath (BS) and vein (V) and between mesophyll (M) and vein (V).

**Supplemental Figure S3**. Enriched transporter families in the six gene expression clusters. **Supplemental Figure S4**. Transcript abundance of genes associated with nitrogen assimilation.

**Supplemental Figure S5**. Transcript abundance of genes associated with the photosynthetic election transport chain (A, B), Calvin Benson Bassham cycle (C), starch biosynthesis (D) and photorespiration (E).

**Supplemental Figure S6**. Transcription factors associated with different clusters. **Supplemental Figure S7**. Transcript abundance of orthogroups shared by bundle sheath cells in rice and Arabidopsis associated with aquaporins, sulphate transport and assimilation, jasmonic acid biosynthesis, and transcription factors.

**Supplemental Figure S8**. Comparison of bundle sheath preferentially expressed genes among different studies.

**Supplemental Figure S9**. Transcript abundance of Aquaporins (*PIP1* and *PIP2*), *ATP sulfurylase* (*ATPS*), *APS reductase* (*APR*), *sulfite reductase* (*SIR*), *13-lipoxygenase* (*LOX*), *allene oxidase cyclase* (*AOC*) and *oxophytodienoate reductase* (*OPR*), *nitrate reductase* (*NIA*) and *nitrite reductase* (*NIR*) in bundle sheath and mesophyll cells of *A. thaliana* (Aubry et al., 2014b), *G. gynandra* (Aubry et al., 2014a), *O. sativa* (this study), *P. virgartum* (Rao et al., 2016), *Z. mays* (Chang et al., 2012), *S. italica* (John et al., 2014) and *S. bicolor* (Emms et al., 2016).

**Supplemental Table S1**. RNA sequencing statistics.

**Supplemental Table S2**. Summary of differential gene expression analysis using DESeq2 and edgeR.

**Supplemental Table S3**. Pairwise comparison and gene expression clusters. **Supplemental Table S4**. Motif enrichment analysis of the six gene expression clusters. **Supplemental Table S5**. Bundle sheath and mesophyll differentially expressed orthologous genes in rice and Arabidopsis.

