## Supplementary material for "The bundle sheath of rice is conditioned to play an active role in water transport as well as sulfur assimilation and jasmonic acid synthesis": S Figure 1

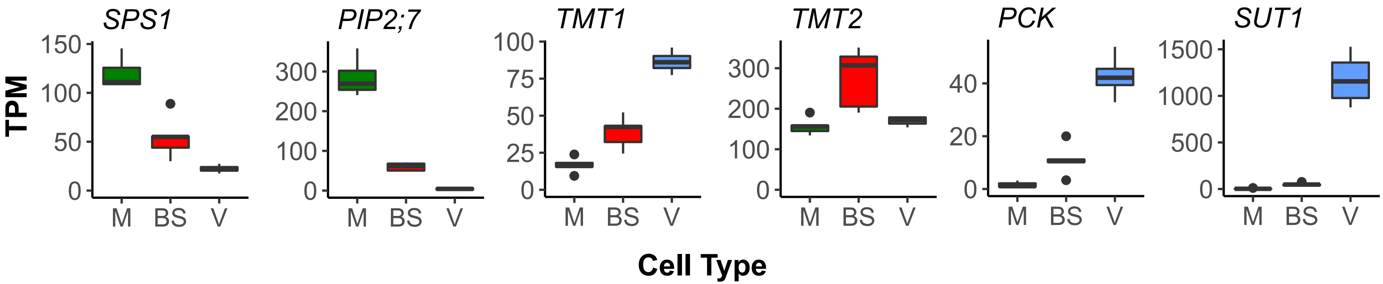


**Supplemental Figure S1. Transcript abundance of genes previously reported to be associated with mesophyll, bundle sheath or veins.** *Sucrose-phosphate synthase 1* (*SPS1*) and *plasma membrane intrinsic protein 2;7* (*PIP2;7*) were preferentially expressed in mesophyll cells. The monosaccharide transporters *TMT1* and *TMT2* were most highly expressed in veins and bundle sheath cells respectively. *Phosphoenolpyruvate carboxykinase* (*PCK*) was most strongly expressed in bundle sheath and veinal cells whilst the *sucrose transporter* (*SUT1*) was most abundant in veins. Data presented as TPM (transcript per million).
