## Supplementary material for "The bundle sheath of rice is conditioned to play an active role in water transport as well as sulfur assimilation and jasmonic acid synthesis": S Figure 2

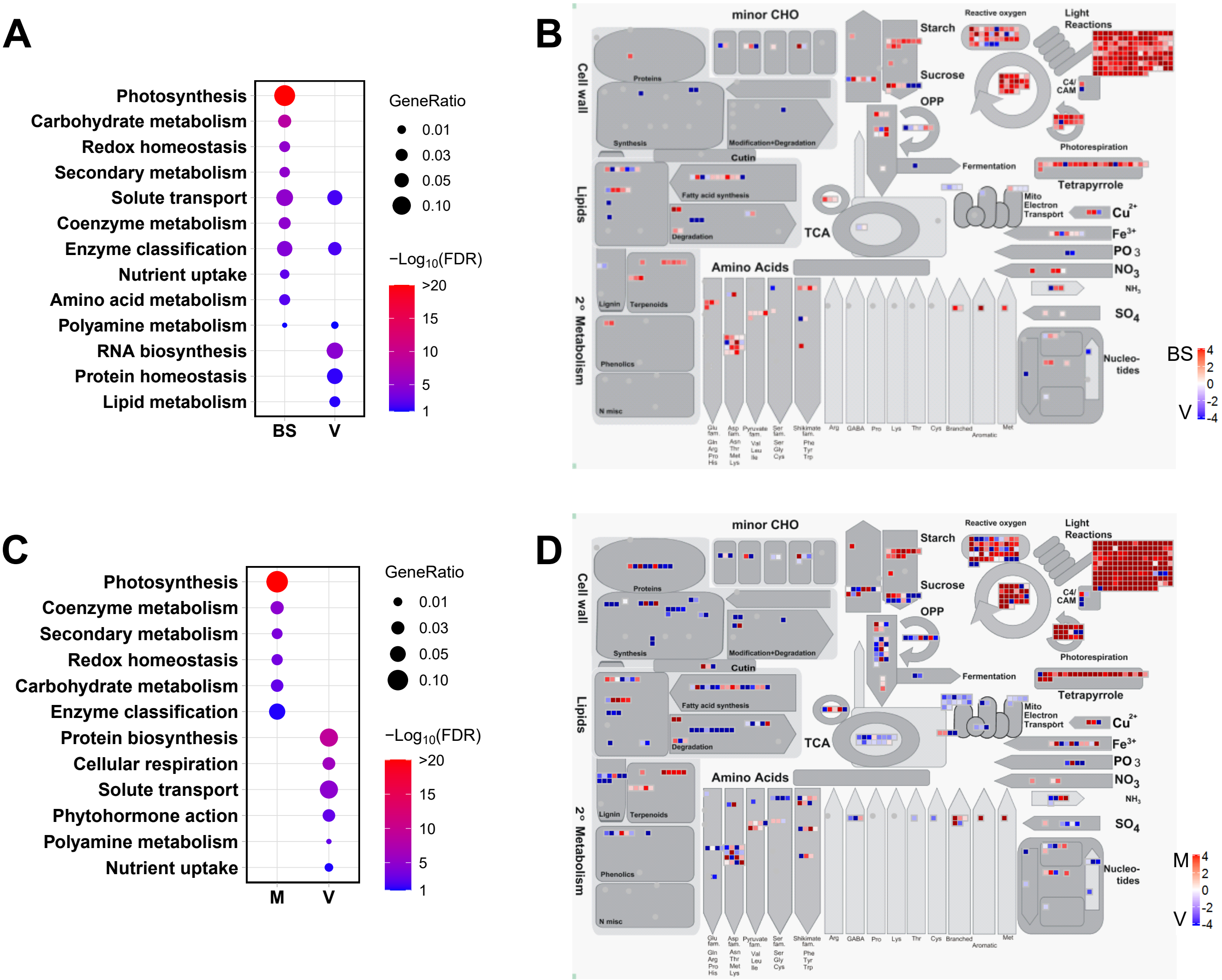


**Supplemental Figure S2. Mapman categories and metabolic overview of pairwise comparisons between bundle sheath (BS) and vein (V) and between mesophyll (M) and vein (V).** (A, C) Primary Mapman categories associated with differentially expressed genes in bundle sheath and veinal cells (A), and mesophyll and veinal cells (C). Terms were defined using Fisher's exact test (False discovery rate, FDR<0.1), colour scale represents negative $\log_{10}$ transformed FDR, gene ratio represents the ratio of the number of matched genes in categories relative to total number of DEGs in each cell type. (B, D). Metabolic overviews of differentially expressed genes between bundle sheath and veins (B) or mesophyll and veins (D). Colour scale represents $\log_2$ fold change.
