## Supplementary material for "The bundle sheath of rice is conditioned to play an active role in water transport as well as sulfur assimilation and jasmonic acid synthesis": S Figure 3

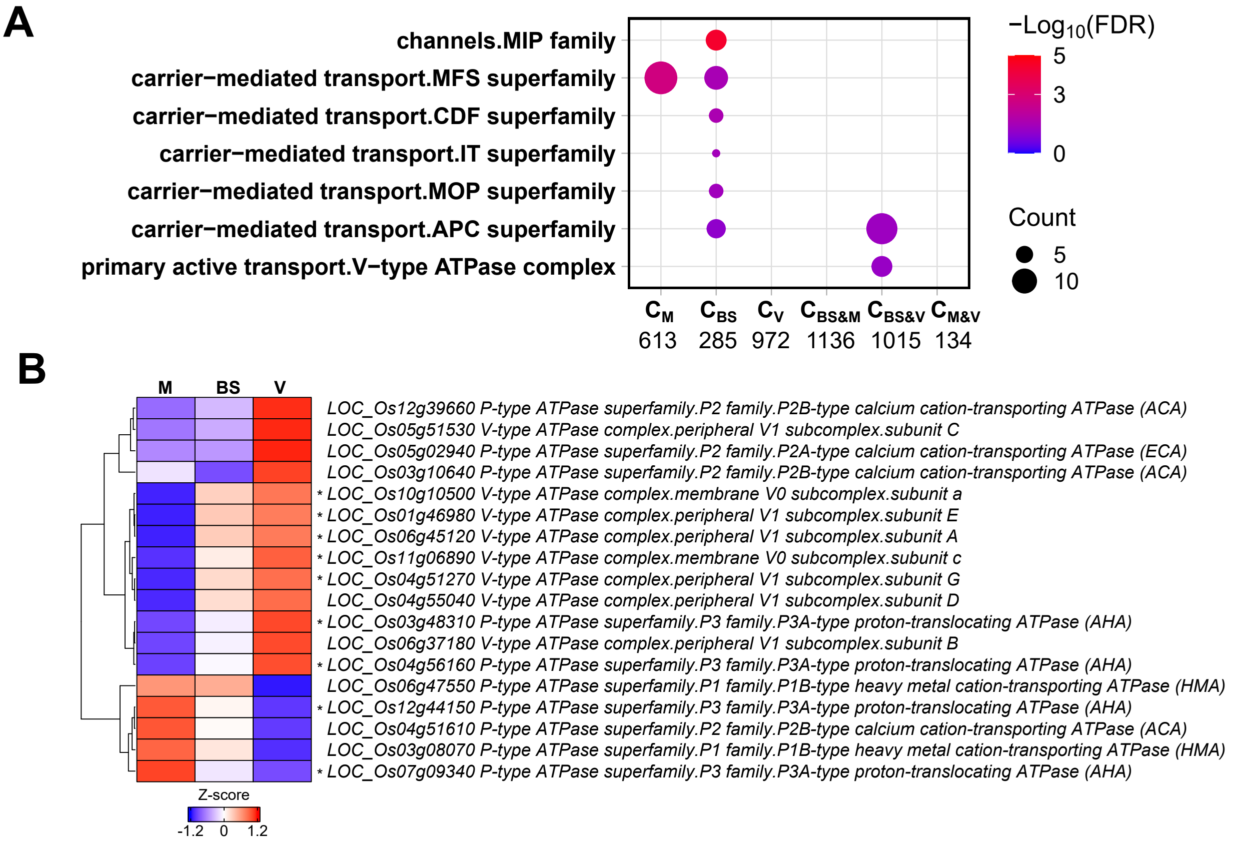


**Supplemental Figure S3. Enriched transporter families in the six gene expression clusters.** (A) Enriched transporter families in the six clusters, colour scale represents negative $\log_{10}$ transformed FDR, significant terms were defined with Fisher's exact test with FDR<0.1. size of dot represents the number of genes found in each category in each cluster. (B) Relative transcript abundance of differentially expressed V and P-type ATPase across M, BS and V, data presented as heatmap of Z-score derived from $\log_2$ transformed quantile normalized TPM. Most V-type ATPase subunits and specific P-type ATPases are more abundant in BS and V, asterisks indicate statistically significant difference between M and BS (FDR and adjust $P < 0.05$ using edgeR and DESeq2 analysis).

Abbreviations for (A): MIP, major intrinsic proteins; CDF, cation diffusion facilitators; MFS, major facilitator superfamily; IT, ion transporter; MOP, multidrug/oligosaccharidyl-lipid/polysaccharide (MOP) flippase superfamily; APC, amino acid-polyamine-organocation superfamily.
