## Supplementary material for "The bundle sheath of rice is conditioned to play an active role in water transport as well as sulfur assimilation and jasmonic acid synthesis": S Figure 4

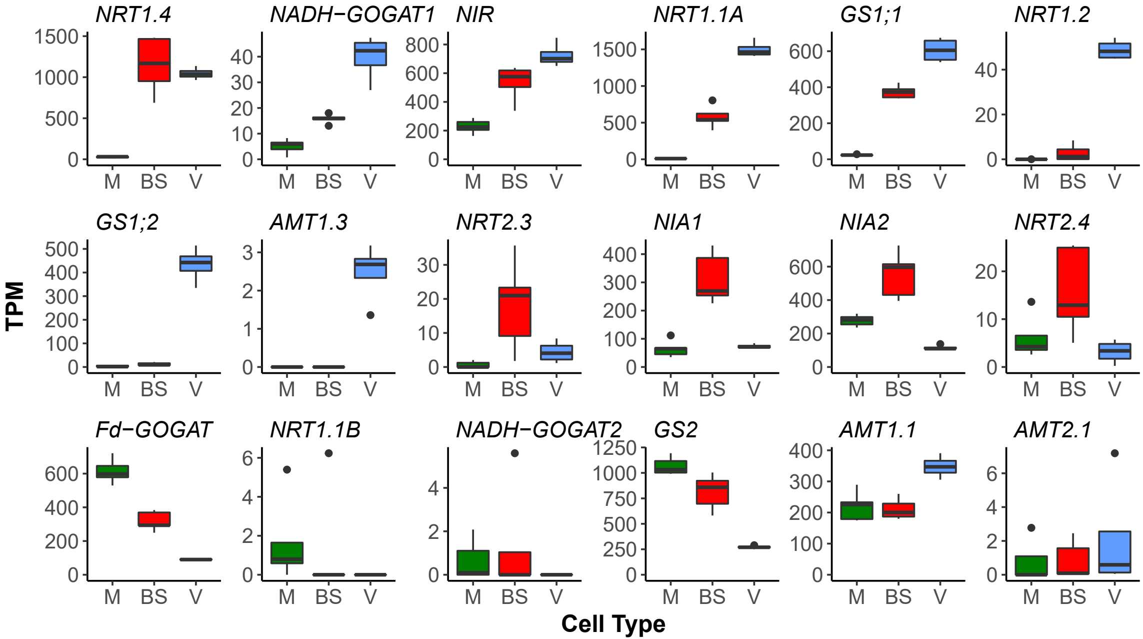


**Supplemental Figure S4. Transcript abundance of genes associated with nitrogen assimilation.** Data presented as TPM (transcript per million).
