## Supplementary material for "The bundle sheath of rice is conditioned to play an active role in water transport as well as sulfur assimilation and jasmonic acid synthesis": S Figure 5

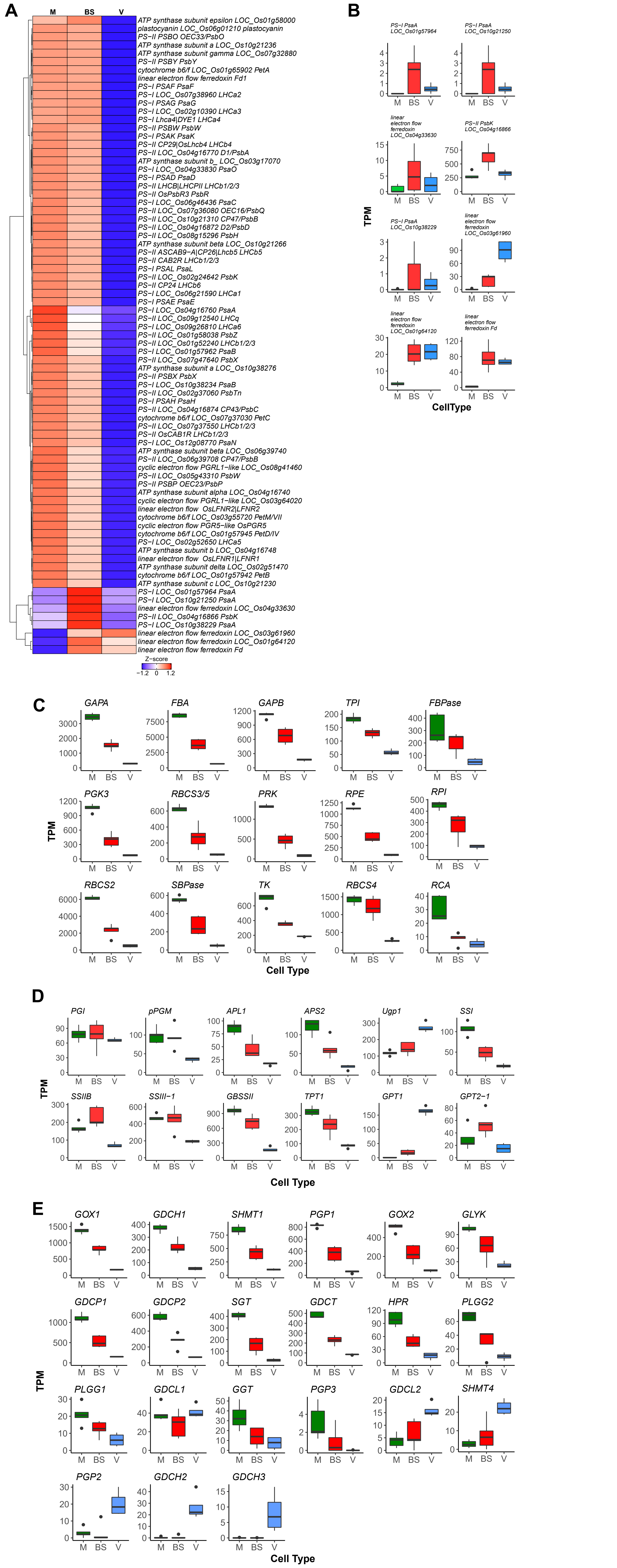


**Supplemental Figure S5. Transcript abundance of genes associated with the photosynthetic election transport chain (A, B), Calvin Benson Bassham cycle (C), starch biosynthesis (D) and photorespiration (E).** Data presented as heatmap of Z-score derived from $log_2$ transformed quantile normalized TPM (A) or TPM (transcript per million) (B-E), gene symbols were extracted from FunRiceGenes database (Yao et al., 2018).

Abbreviations in (A, B): PSII, photosystem II; PSI, photosystem I; LHC, light-harvesting complex; FLFNR, leaf-type ferredoxin-NADP oxidoreductase; PGR, proton gradient regulation. (D): PGI, plastidial phosphoglucose isomerase; PGM, plastidial phosphoglucomutase; APL1, ADP-glucose pyrophosphorylase; APS2, ADP-glucose pyrophosphorylase; Ugp1, cytosolic UDP-glucose pyrophosphorylase; SS, starch synthase; GBSS, granule-bound starch (amylose) synthase; TPT, triose phosphate/ phosphate translocator; GPT, glucose 6-phosphate/ phosphate translocator.
