## Supplementary material for "The bundle sheath of rice is conditioned to play an active role in water transport as well as sulfur assimilation and jasmonic acid synthesis": S Figure 6

**A**

| Family | $TF_M$ | $TF_{BS}$ | $TF_V$ | $TF_{BS\&M}$ | $TF_{BS\&V}$ | $TF_{M\&V}$ |
|---|---|---|---|---|---|---|
| ERF | 3 | 2 | 4 | 0 | 4 | 1 |
| HD-ZIP | 3 | 1 | 3 | 0 | 3 | 0 |
| MYB_related | 3 | 0 | 5 | 2 | 3 | 3 |
| bZIP | 2 | 1 | 7 | 0 | 4 | 0 |
| bHLH | 2 | 1 | 6 | 2 | 3 | 1 |
| CO-like | 2 | 0 | 2 | 6 | 1 | 0 |
| C2H2 | 2 | 0 | 2 | 4 | 2 | 0 |
| MYB | 0 | 1 | 2 | 0 | 6 | 0 |
| G2-like | 0 | 0 | 6 | 3 | 4 | 0 |
| Dof | 0 | 0 | 4 | 1 | 3 | 0 |
| DBB | 2 | 0 | 2 | 1 | 0 | 0 |
| NAC | 1 | 1 | 3 | 1 | 2 | 0 |
| WRKY | 1 | 1 | 2 | 0 | 4 | 0 |
| ARF | 0 | 0 | 2 | 0 | 1 | 0 |
| MIKC_MADS | 0 | 0 | 2 | 0 | 1 | 0 |
| GRAS | 1 | 1 | 1 | 1 | 2 | 0 |
| TALE | 1 | 1 | 1 | 0 | 0 | 0 |
| ZF-HD | 1 | 0 | 3 | 0 | 0 | 1 |
| C3H | 1 | 0 | 1 | 1 | 1 | 0 |
| Nin-like | 1 | 0 | 1 | 1 | 0 | 0 |
| LSD | 1 | 0 | 1 | 0 | 0 | 0 |
| GATA | 1 | 0 | 0 | 2 | 1 | 2 |
| Trihelix | 1 | 0 | 0 | 1 | 1 | 0 |
| CAMTA | 1 | 0 | 0 | 0 | 0 | 1 |
| EIL | 1 | 0 | 0 | 0 | 0 | 0 |
| HB-other | 1 | 0 | 0 | 0 | 0 | 0 |
| HSF | 1 | 0 | 0 | 0 | 0 | 0 |
| NF-YC | 1 | 0 | 0 | 0 | 0 | 0 |
| NF-YB | 0 | 0 | 1 | 0 | 1 | 0 |
| GRF | 0 | 0 | 1 | 0 | 0 | 1 |
| ARR-B | 0 | 0 | 1 | 0 | 0 | 0 |
| B3 | 0 | 0 | 1 | 0 | 0 | 0 |
| GeBP | 0 | 0 | 1 | 0 | 0 | 0 |
| SBP | 0 | 0 | 1 | 0 | 0 | 0 |
| AP2 | 0 | 0 | 0 | 1 | 0 | 0 |
| LBD | 0 | 0 | 0 | 1 | 0 | 0 |
| FAR1 | 0 | 0 | 0 | 0 | 1 | 1 |
| BES1 | 0 | 0 | 0 | 0 | 1 | 0 |
| HRT-like | 0 | 0 | 0 | 0 | 1 | 0 |
| M-type_MADS | 0 | 0 | 0 | 0 | 1 | 0 |
| VOZ | 0 | 0 | 0 | 0 | 1 | 0 |

**B**

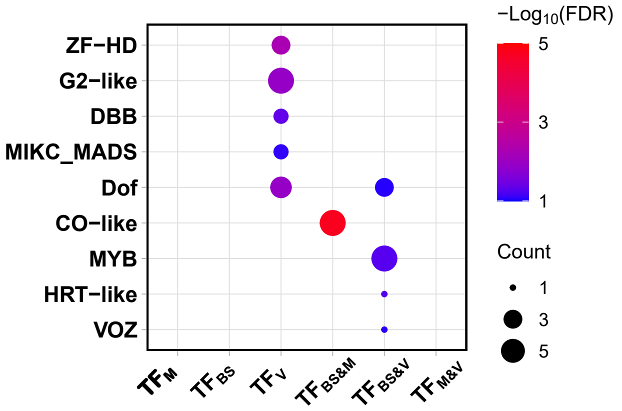


**C**

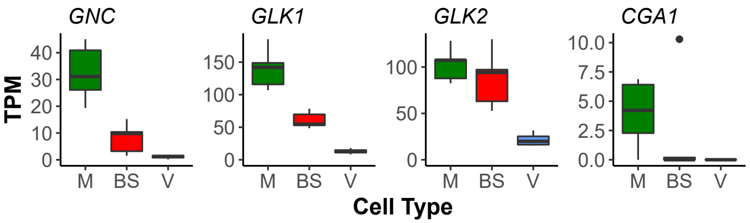


**Supplemental Figure S6. Transcription factors associated with different clusters.** (A) Number of transcription factors in each families across the six gene expression clusters, which is also illustrated in scale of red. (B) Over-represented transcription factor families defined by Fisher's exact test (FDR<0.1). (C) transcript abundance of known transcription factors regulating chloroplast biogenesis, data presented in TPM (transcript per million).
