## Supplementary material for "The bundle sheath of rice is conditioned to play an active role in water transport as well as sulfur assimilation and jasmonic acid synthesis": S Figure 7

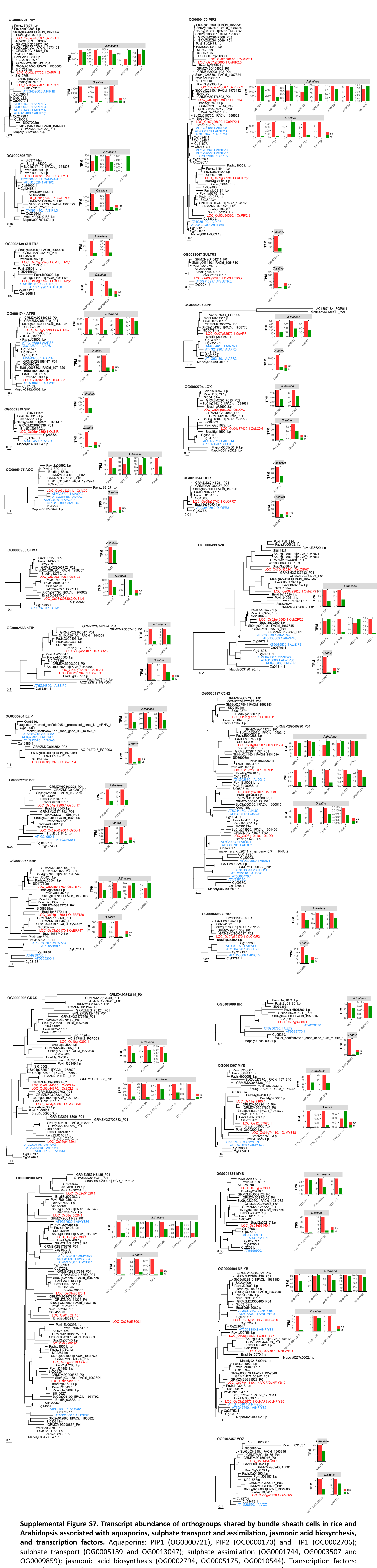

**Supplemental Figure S7. Transcript abundance of orthogroups shared by bundle sheath cells in rice and Arabidopsis associated with aquaporins, sulphate transport and assimilation, jasmonic acid biosynthesis, and transcription factors.** Aquaporins: PIP1 (OG0000721), PIP2 (OG0000170) and TIP1 (OG0002706); sulphate transport (OG0005139 and OG0013047); sulphate assimilation (OG0001744, OG0003507 and OG0009859); jasmonic acid biosynthesis (OG0002794, OG0005175, OG0010544). Transcription factors: SLIM1 (OG0003985), Basic Leucine Zipper (OG0000499, OG0002563, OG0005764), $C_2H_2$-type Zinc Finger (OG0000197), DNA-binding with One Finger (OG0002717), Ethylene Responsive Factor (OG0000957), GRAS (OG0000296, OG0005593), Hairy-Related Transcription-Factor (OG0005688), MYB (OG0000100, OG0001367 and OG0001681), Nuclear Factor-YB (OG0000404), and Vascular Plant One-Zinc Finger Protein (OG0002457). Data presented as the mean of TPM, rice and Arabidopsis loci are highlighted with red and blue text respectively, asterisks indicate statistically significant difference (FDR and adjust $P < 0.05$ using edgeR and DESeq2 analysis in this study, PPDE>0.95 in Aubry et al., 2014b).
