## Supplementary material for "The bundle sheath of rice is conditioned to play an active role in water transport as well as sulfur assimilation and jasmonic acid synthesis": S Figure 8

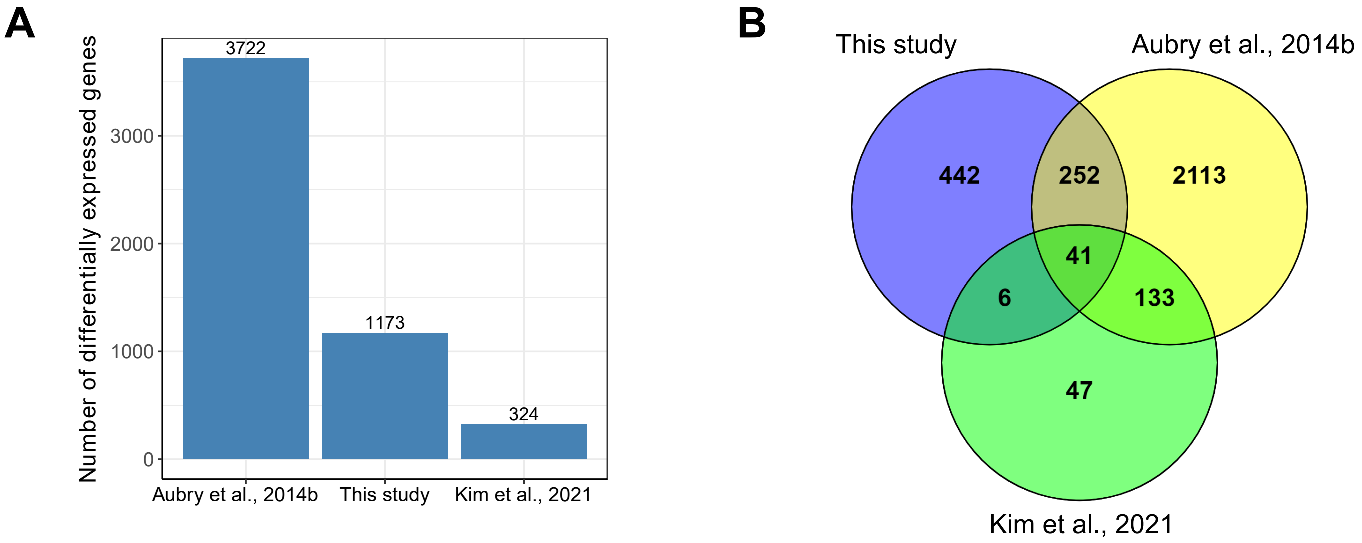


**Supplemental Figure S8. Comparison of bundle sheath preferentially expressed genes among different studies.** (A) Number of bundle sheath preferentially expressed genes identified in Aubry et al., 2014b, Kim et al., 2021 and this study. (B) Overlapping of orthogroups containing bundle sheath differentially expressed genes between rice (this study) and Aubry et al., 2014b, Kim et al., 2021 respectively.
