## Supplementary material for "The bundle sheath of rice is conditioned to play an active role in water transport as well as sulfur assimilation and jasmonic acid synthesis": S Figure 9

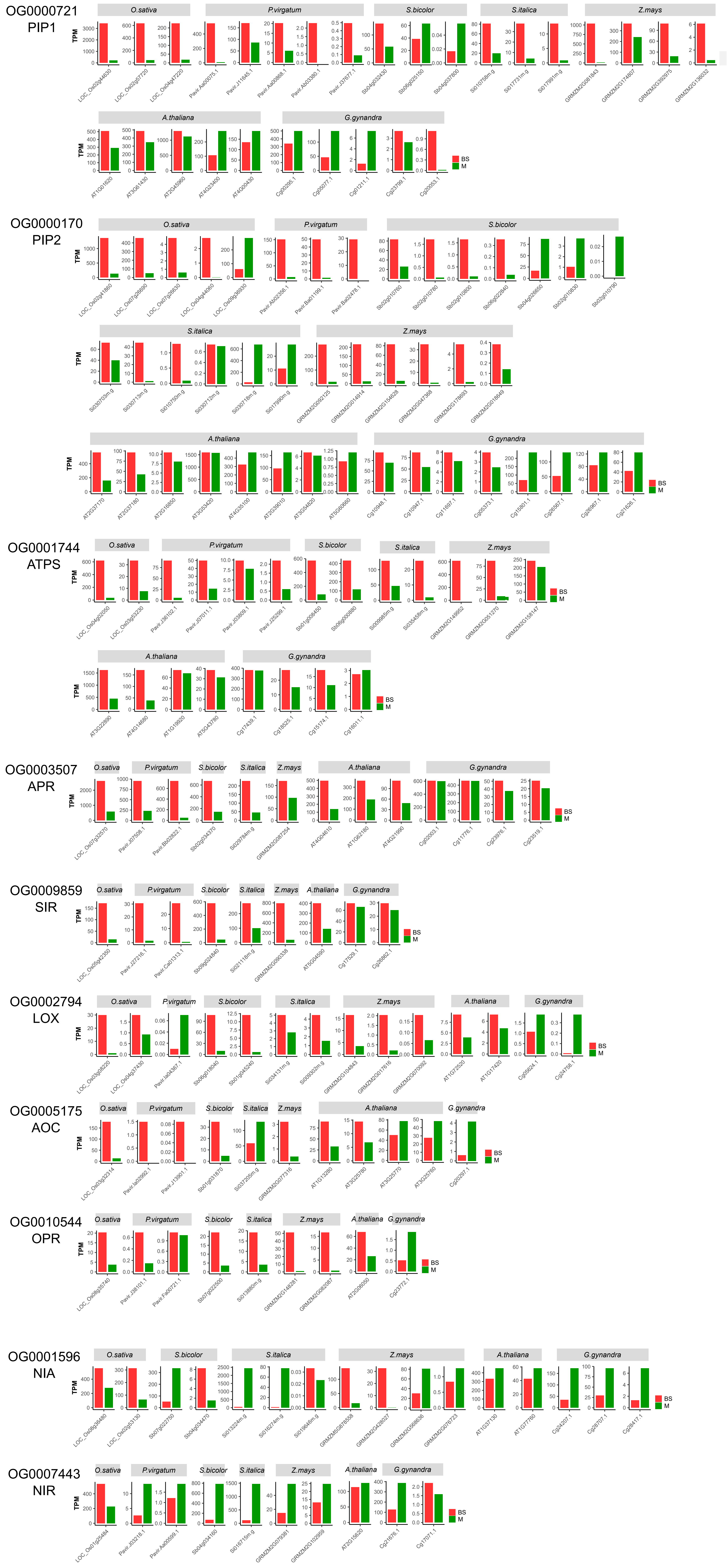


**Supplemental Figure S9. Transcript abundance of aquaporins (*PIP1* and *PIP2*), *ATP sulfurylase* (*ATPS*), *APS reductase* (*APR*), *sulfite reductase* (*SIR*), *13-lipoxygenase* (*LOX*), *allene oxidase cyclase* (*AOC*) and *oxophytodienoate reductase* (*OPR*), *nitrate reductase* (*NIA*) and *nitrite reductase* (*NIR*) in bundle sheath and mesophyll cells of *A. thaliana* (Aubry et al., 2014b), *G. gynandra* (Aubry et al., 2014a), *O. sativa* (this study), *P. virgatum* (Rao et al., 2016), *Z. mays* (Chang et al., 2012), *S. italica* (John et al., 2014) and *S. bicolor* (Emms et al., 2016).** Data presented as mean of TPM.
